# Impact of Trastuzumab and Pertuzumab on HER2 Localization and the Estrogen Receptor Cistrome in ER+/HER2+ Breast Cancer

**DOI:** 10.64898/2026.09.07.749891

**Authors:** Stanley Tam, Shaymaa Bahnassy, Lu Jin, Matthew D. McCoy, Suman Ranjit, Raneen Rahhal, Alec T. McIntosh, Douglas K. Kung, Gong Tang, Sandra M. Swain, Rebecca B. Riggins

## Abstract

**Background:** Although anti-HER2 monoclonal antibody therapy with trastuzumab and pertuzumab is highly effective for HER2-positive breast cancer, co-expression of estrogen receptor (ER) significantly reduces pathologic complete response rates.

**Objective:** We investigated how HER2-targeted inhibition affects HER2 cellular localization and ER genomic binding in ER-positive, HER2-positive breast cancer.

**Methods:** We evaluated HER2 localization in a tissue microarray of treatment-naive patients. Using cellular fractionation, immunofluorescence, chromatin immunoprecipitation, and genome-wide profiling (CUT&RUN), we evaluated HER2 localization and ER genomic binding following acute anti-HER2 treatment and in models of treatment resistance. The treatment-induced ER-bound gene signature was assessed for associations with pathologic complete response and survival in breast cancer clinical cohorts.

**Results:** In treatment-naive primary breast tumors, nuclear HER2 inversely correlated with ER levels. In cell models, treatment with trastuzumab and pertuzumab induced nuclear and chromatin accumulation of HER2. Concurrently, acute anti-HER2 treatment displaced ER from canonical estrogen response elements at classical target genes, yet genome-wide profiling revealed redistribution of ER binding toward non-canonical zinc finger motifs adjacent to pro-survival Wnt and RAGE pathway genes (*FZD8*, *RELA*). Expression of a drug-induced ER-bound gene signature was significantly higher in tumors of individuals who did not achieve pathologic complete response following neoadjuvant anti-HER2 therapy and significantly correlated with reduced distant metastasis-free survival in clinical cohorts.

**Conclusions:** Anti-HER2 targeted therapy causes dynamic cistromic reprogramming of ER from classical estrogen pathways toward alternative pro-survival networks. These findings implicate ER redistribution as a possible mediator of resistance and highlight novel therapeutic targets in ER-positive, HER2-positive breast cancer.

## Introduction

Breast cancer is a major contributor to the cancer-related healthcare burden in the US, accounting for an estimated 321,000 new cases in 2026, according to the National Cancer Institute SEER cancer epidemiology program (1). Breast cancer is commonly divided into subtypes based on the expression of several receptors that guide treatment decisions (2–4), including the hormone receptor (HR) estrogen receptor alpha (ER) and human epidermal growth factor receptor 2 (HER2). ER+ breast cancer is treated with endocrine therapies that inhibit the ER pathway (5,6), whereas the HER2+ subtype is treated with anti-HER2 therapies that either inhibit the receptor directly or use antibody-drug conjugates to deliver cytotoxic chemotherapies specifically to cancer cells (7).

The ER+/HER2+ subtype is characterized by the expression of ER and the overexpression of HER2 and/or amplification of the gene that encodes HER2, *ERBB2* (8,9). While there are multiple therapeutic options for treating ER+/HER2+ breast cancer, the type and sequencing of these treatments can vary from patient to patient. The American Society of Clinical Oncology guidelines recommend treating patients with HER2+ breast cancer in the neoadjuvant setting with chemotherapy plus anti-HER2 therapy, such as the combination of monoclonal antibodies trastuzumab and pertuzumab (TP) as first-line therapy, regardless of ER expression (10). Endocrine therapy and anti- HER2 therapies can be used as monotherapy or in various combinations, and TP is one of the most effective neoadjuvant treatments for HR+/HER2+ disease (11,12). However, expression of one or more HRs is associated with reduced pathologic complete response (pCR) following neoadjuvant anti-HER2 therapy compared to HR-/HER2+ breast cancer (13–18). Similarly, the expression of HRs is also associated with reduced effectiveness of anti-HER2 therapies in patients with metastatic disease (19). Additionally, higher expression of ER-related genes is associated with lower pCR in the NA-PHER2 clinical trial among patients receiving TP with or without fulvestrant (20).

These clinical observations highlight the need to define simultaneous contributions of both ER and HER2 to anti-HER2 targeted therapy resistance. ER may promote tumor survival independently, or act as a compensatory mechanism when HER2 is inhibited. Preclinical cell line models of ER+/HER2+ breast cancer offer a useful setting to identify mechanisms underlying ER-mediated reduction of anti-HER2 therapy efficacy, and to test whether anti-HER2 treatment itself contributes to these mechanisms. In this study, we investigated the localization and functions of ER and HER2 in response to anti-HER2 therapy in ER+/HER2+ breast cancer.

We show that in primary treatment-naive ER+/HER2+ breast tumors, high nuclear HER2 is associated with reduced ER levels and that TP induces HER2 nuclear and chromatin localization in ER+/HER2+ breast cancer cell line models. TP reduces ER binding to regulatory sites of several well-established canonical ER target genes, but induces ER binding at other genes related to Wnt signaling and receptor for advanced glycation endproducts (RAGE) signaling. Importantly, expression of several of these TP-induced ER target genes is associated with shorter survival or reduced likelihood of achieving pCR with neoadjuvant anti-HER2 treatment. Collectively, these findings highlight ER redistribution following anti-HER2 therapy as a mediator of resistance and suggest candidate therapeutic targets in ER+/HER2+ breast cancer.

## Materials and Methods

### Cell culture and treatments

BT-474 (RRID:CVCL_0179), MDA-MB-361 (RRID:CVCL_0620), MCF-7 (RRID:CVCL_0031), and SK-BR-3 (RRID:CVCL_0033) breast cancer cells were obtained from the Georgetown University Tissue Culture and Biobanking Shared Resource. BT-474 trastuzumab/pertuzumab-resistant cells (BT-474 TP resistant) were generated by continuously culturing parental cells with trastuzumab (T; Genentech, or MedChemExpress, Cat# HY-P9907, RRID:AB_2938985) and pertuzumab (P; Genentech, or MedChemExpress Cat# HY-P9912, RRID:AB_3694970), starting at 0.1 µg/ml and increasing by 0.1 µg/ml every 2 passages over several months until 1.5 µg/ml was reached. BT-474 TP resistant cells were subsequently maintained in 1.5 µg/ml TP, except for when cells were plated the day before most experiments in TP-free media. All cell lines were cultured in IMEM (Thermo Cat# A1048901) supplemented with 10% fetal bovine serum, maintained at 37°C in a humidified incubator with 5% CO_2_, and routinely tested for *Mycoplasma* contamination.

Unless otherwise indicated, cells were treated with, 100 nM fulvestrant (Selleckchem Cat# S1191), 100 µg/ml T and/or P, 100 nM tucatinib (MedChemExpress Cat# HY-16069), or 100 ng/ml epidermal growth factor (EGF; Thermo Cat# PHG0311) for 3 hours. For RNA expression experiments, treatments were extended to 24 hours. For hormone-deprivation experiments (RNA-seq, ChIP-PCR, ChIP-seq), cells were cultured in phenol red-free MEM media supplemented with 10% charcoal-stripped FBS for 3 days then stimulated with 10 nM 17 beta-estradiol (E2, Sigma Cat# E2758) for 3 hours.

### Western blot analysis of whole-cell and fractioned lysates

Cells were plated and treated the following day as indicated for 3 hours, collected by scraping, and washed twice with cold PBS. Whole-cell lysates were prepared in RIPA buffer (150 mM NaCl, 50 mM Tris-HCl, 1% IGEPAL CA-630 - Sigma Cat# I8896, 0.5% sodium deoxycholate, Roche cOmplete protease inhibitor cocktail - Sigma Cat# 04693116001).

For subcellular fractionation, cells were lysed, and the cytoplasmic fractions were isolated by sucrose density centrifugation as described by Li et al. (21). The resulting nuclei were washed and lysed in cold nuclear lysis buffer (0.1 M NaCl, 0.7% Triton X-100, 40% sucrose). Chromatin pellets were collected by centrifugation and sonicated using a Bioruptor Pico sonicator (Diagenode) for 10 seconds on, 30 seconds off for ∼8 cycles until the solution became clear.

All lysates were denatured by boiling at 95°C, with 4X LDS loading dye (Thermo Cat# NP0008) and beta-mercaptoethanol (Sigma Cat# M3148), resolved on 4-12% Bis-Tris gradient gels (Thermo Cat# NP0321), and transferred to nitrocellulose membranes. Membranes were blocked for 1 hour in 1% milk in tris-buffered saline with Tween-20 (Sigma Cat# P1379), incubated overnight with primary antibodies, followed by a 1 hour incubation with appropriate secondary antibodies (Table S1), washed, developed using chemiluminescent substrate, and imaged using Amersham Imager 600 (GE Healthcare Life Sciences).

### Immunofluorescence microscopy

BT-474 TP resistant cells were maintained in 1.5 µg/ml TP or cultured without TP for 72 hours. Both parental and TP resistant cells were seeded (70,000 cells/well) onto poly-L-lysine (Sigma Cat# P6282) coated coverslips in 12-well plates. After 48 hours, only parental cells were treated with vehicle or 100 µg/ml TP for 3 hours. All coverslips were washed with PBS, fixed in 4% paraformaldehyde, permeabilized with 0.02% Triton X-100 (Sigma Cat# X100) for 10 mins, and blocked in a 10% goat serum solution (Thermo Cat# 50062Z). Samples were incubated overnight at 4 °C with anti-HER2 monoclonal antibody (1:150), washed, and incubated with Alexa Fluor 594- conjugated secondary antibody (1:400) for 1 hour at room temperature (antibodies listed in Table S1). Nuclei were counterstained with DAPI (Thermo Cat# D1306), and coverslips mounted with Fluoro-Gel mounting medium (Electron Microscopy Sciences Cat# 17985-30).

High resolution imaging was performed using spinning disk confocal microscopy on an inverted Nikon Ti2-E microscope (CSU-W1 SoRa; Nikon Instruments) equipped with 405 nm and 561 nm wavelength lasers and a 100X NA 1.45 oil immersion objective. Z-stacks were acquired using a NIDAQ Piezo Z Drive with a step size of 0.06 µm, spanning a total range of 0.72 µm around the focal center (13 planes in total). Using NIS Elements (v5.41.01; Nikon), the artificial intelligence denoise plugin and 4 iterations of Richardson-Lucy deconvolution were applied to maximum intensity Z-projections of confocal images acquired under identical laser settings. Images were analyzed in Fiji (NIH), and nuclear HER2 fluorescence was quantified as Integrated Density (22).

### Crystal violet growth assay

Cells were plated in 96-well plates at 10,000 cells/well for BT-474 and BT-474 TP resistant cells, and 30,000 cells/well for MDA-MB-361. The next day, treatments were initiated and drug-containing media was replenished every 3 days. After 8 days, cells were stained with 0.5% crystal violet in 25% methanol solution, washed and quantified using a BioTek ELx808 plate reader measuring absorbance at 550 nm with background correction at 405 nm.

### Quantitative PCR (qPCR)

For mRNA quantifications, cells were treated for 24 hours, scraped, and washed in cold PBS. RNA was extracted using a PureLink RNA mini kit (Thermo Cat# K310001), and cDNA was prepared using iScript cDNA synthesis kit (Bio-Rad Cat# 1708891). Gene expression was quantified by real-time PCR using the iTaq SYBR green mix (Bio-Rad) with gene-specific primers (Table S2), normalized to beta-actin or beta-tubulin, and analyzed with the ΔΔCT method.

### Chromatin immunoprecipitation (ChIP), ChIP-qPCR and ChIP sequencing (ChIP-seq)

Cells were treated with anti-ER or anti-HER2 therapies for 3 hours, fixed in 1% formaldehyde for 15 minutes at room temperature, and quenched with glycine. In the case of E2 stimulation, cells were hormone-deprived and treated with E2 for 3 hours as described above, prior to fixation.

For ChIP-PCR, chromatin was isolated using a Cell Signaling Sonication ChIP kit (Cat# 56383) and sonicated for 30 seconds on/off for 6 cycles. ChIP was performed using anti-ER alpha, anti-HER2 or IgG antibodies (Table S1). DNA was purified with a DNA PCR purification kit (Qiagen Cat# 28104) and qPCR was then performed using Bio-Rad iTaq SYBR green mix (Bio-Rad Cat# 1725120) and primers (Table S2). Data were presented as fold enrichment relative to input.

For ER ChIP-seq, fixed cells were snap-frozen and sent to Active Motif for processing. Samples were lysed with a Dounce homogenizer, and chromatin was sonicated to 300-500 bp. An aliquot was reserved as input, and the remaining chromatin was immunoprecipitated with anti-ER alpha antibody (Table S1). Samples were treated with RNase and proteinase K, de-crosslinked by heating, and purified by phenol-chloroform extraction. Libraries were prepared as described (23), amplified by PCR, and sequenced by Illumina NovaSeq to produce 75 nucleotide long single-end reads. Reads were aligned to the hg38 Human Reference Assembly with the default configuration of BWA, peaks were called with MACS2 (24,25), and Bedtools intersect was used to determine differentially enriched peaks across experimental conditions (26). For the heatmaps, ChIP-seq signal was normalized to the input signal and visualized with deepTools (27). To predict ER-bound target genes with E2-modulated expression and associated enriched motifs, we used BETA (28), integrating parallel ER ChIP-seq and RNA-seq data from BT-474 cells alongside previously published MCF-7 datasets (29).

### RNA Sequencing (RNA-seq)

BT-474 cells were plated in 10 cm dishes, hormone-deprived as described above, and treated with 10 nM E2 for 3 hours. Cells were scraped, snap-frozen, and sent to Active Motif for processing. Total RNA was isolated using the Qiagen RNeasy Mini Kit (Qiagen Cat# 74104). Libraries were prepared from 2 µg of RNA with Illumina’s TruSeq Stranded mRNA Library kit (Cat# 20020594) and sequenced on NextSeq 500 as paired-end 42-nt reads. Sequence reads were analyzed with the STAR alignment - DESeq2 software pipeline. Gene set enrichment analysis (GSEA) was performed using MSigDB v2026.1.Hs accessed January 2026 using the Reactome, Gene Ontology (GO), and Hallmark geneset databases (30,31) on the genes identified by BETA.

### Cleavage Under Targets & Release Using Nuclease (CUT&RUN)

BT-474 cells were plated and treated the next day with 100 µg/ml TP for 3 hours. Nuclei were isolated from native (non-fixed) cells using a buffer (EpiCypher Cat# 21-1026), snap-frozen, and sent to EpiCypher. CUTANA™ CUT&RUN was performed using the autoCUT&RUN protocol, derived from previously described studies (32–34). In brief, for each CUT&RUN reaction, 500K nuclei were dispensed to individual wells of a 96-well plate, immobilized onto Concanavalin-A beads (Con-A; EpiCypher Cat# 21-1401), and incubated overnight at 4°C with 0.5 µg ER alpha or control (IgG, H3K4me3, H3K27me3) antibodies (Table S1). pAG-MNase (EpiCypher Cat# 15-1016) was added for 2 hours at 4°C. Reactions were stopped with Stop buffer (EpiCypher Cat# 21-1003k) supplemented with 0.5 ng of *E. coli* DNA (EpiCypher Cat# 18-1401k). Enriched DNA was purified with Serapure beads using a 2:1 bead-to-DNA ratio. Recovered DNA was quantified with PicoGreen, normalized to 5 ng DNA before preparing sequencing libraries (CUTANA CUT&RUN Library Prep kit; EpiCypher Cat# 14-1001). All autoCUT&RUN steps were performed on Tecan Freedom EVO robotics platforms, and libraries were sequenced on Illumina NextSeq2000, 2x50bp paired-end reads.

Raw sequencing reads were quality-assessed with FastQC (v0.11.9), trimmed of adapter sequences and low-quality bases with Cutadapt (v3.5), and filtered to retain reads longer than 25 bp. Trimmed paired-end reads were aligned to the human reference genome (GRCh38; Ensembl release 105) with Bowtie2 (v2.5.0); only properly paired and uniquely mapped reads were retained. Peaks were called with MACS2 (v2.2.7.1) with default settings for narrow peaks, and signal intensity was quantified using normalized MACS2 peak scores. Motif enrichment was analyzed with HOMER (v5.1), and peaks were assigned to genes by proximity to transcription start sites. Genes were categorized as upregulated or downregulated based on the direction and magnitude of peak score differences between treated and control conditions. CUT&RUN tracks were visualized in the IGV-Web app v2.4.3 (35).

### Multiplex immunohistochemistry (IHC) for HER2+ tissue microarray (TMA)

The HER2+ breast invasive ductal carcinoma TMA was constructed by the Histopathology and Tissue Shared Resource (HTSR) at Georgetown University Medical Center Lombardi Comprehensive Cancer Center as described previously (36). Patients provided informed consent to the HTSR, the Survey, Recruitment, and Biospecimen Shared Resource at Lombardi, and/or Indivumed under the respective GUMC protocols: 1992-048, Pr0000007, and 2007-345. Of the 50 HER2+ cases, 31 were HR+/HER2+ and 19 were HR-/HER2+. Demographic, clinical and pathological characteristics were summarized previously (36).

Sections of 5 μm thickness were cut from FFPE tissue blocks containing donor or recipient tissue samples or tonsil control tissue blocks. The slides were baked at 60 °C, deparaffinized in xylene, rehydrated, washed in tap water and incubated with 10% neutral buffered formalin (NBF) for an additional 20 minutes to increase tissue-slide retention. Epitope retrieval/microwave treatment (MWT) for all antibodies was performed by boiling slides in Antigen Retrieval buffer 6 (AR9 pH9; Akoya, AR9001KT). Protein blocking was performed using antibody diluent/blocking buffer (Akoya, ARD1001EA) for 10 minutes at room temperature. Primary antibody/OPAL dye pairings, staining order and incubation conditions for the antibodies used for multiplex IHC are listed in Table S3. DAPI was used as a nuclear counterstain. Multispectral images were acquired by the Vectra3 multispectral imaging platform (Akoya Biosciences) and processed with Phenochart and inForm 2.4.6 (PerkinElmer/Akoya) as described previously (37,38).

Total and nuclear HER2 expression were quantified in individual tumor cells (identified as pan-cytokeratin-positive cells, panCK+) and nuclei based on immunofluorescence intensity, with nuclear HER2 defined by the overlap with DAPI-stained regions. Coordinates were assigned to TMA cores, and a custom MATLAB script extracted coordinate-specific intensity values for individual cells and nuclei within each core. It then calculates the average expression per cell and the standard deviation for each sample associated with the coordinate IDs. These values are subsequently mapped to grid identifiers and then to randomly generated patient IDs to obtain individual cell expression for each patient. Total (whole cell) and nuclear HER2 mean fluorescence intensity (MFI) were compared between ER-/HER2+ and ER+/HER2+ cases.

### Use and Re-Use of Publicly Available or Previously Published Data

METABRIC (39–41) data were accessed via KMPlotter (42) and cBioPortal (43–45), and the pooled dataset of n=123 HER2+ breast cancers treated with anti-HER2 therapy were accessed via ROCPlot (46), on June 28, 2025. For the NSABP B-41 clinical trial, RNAseq count data originally published by Fernandez-Martinez et al. were normalized with Salmon, which performs quasi-mapping-based transcript quantification (47). The length-scaled counts were then log-transformed with base 2 for the downstream differential expression analyses. For each putative ER target, a two sample t-test was used to compare the expression level by pCR status overall, and separately in the ER- and ER+ subgroups. Both raw p-values and the q-values, from the Benjamini-Hochberg procedure that controls the false discovery rate, are reported.

## Statistical analysis

All bar charts were plotted as mean ± standard deviation. For data visualization of bar charts of RNA expression, and crystal violet growth assays, the data are shown as the average of 3 independent biological replicates. For visualization of ChIP-qPCR bar charts, data are represented as 1 representative biological replicate out of 3 independent replicates. For visualization of immunofluorescence quantification scatter plots, the data represents 2 independent replicates.

Unpaired t-test or one-way ANOVA with Dunnett’s multiple comparisons were used to calculate statistical significance in most experiments, except as noted in the figure legends. Statistical analyses were performed in GraphPad Prism (v10.0.3), with p < 0.05 considered significant.

## Results

### ER and HER2 are negatively correlated in ER+/HER2+ breast cancer

ER+/HER2+ breast tumors can display intratumoral heterogeneous expression of ER and HER2 (48,49). We determined ER and HER2 expression patterns using multiplex IHC staining in a tissue microarray (TMA) of 50 HER2+ (HER2-amplified) treatment-naive primary breast cancers: 31 ER+ (>10% positive cells by diagnostic pathology) and 19 ER- tumors (Fig. 1A-1C). We first characterized ER and HER2 expression to identify associations between their expression patterns. Total HER2 mean fluorescence intensity (MFI) is not significantly different between ER+ and ER- tumors, however, there is a negative correlation between HER2 MFI and percent ER positive cells (Fig. 1B). This relationship prompted us to examine the distribution within the nuclear cell compartment, and surprisingly, we observed that several tumors had HER2 staining in the nucleus. When we considered the ER status of the tumors, we observed that ER- tumors had higher nuclear HER2 intensity compared to ER+ tumors and a trend toward a negative correlation between nuclear HER2 MFI and percent ER positive cells, although this correlation was not statistically significant (Fig. 1C). Together, these data demonstrate an inverse correlation between HER2 expression and ER status or percent positivity, and suggest that ER and HER2 nuclear localization may be linked.

**Figure 1.**
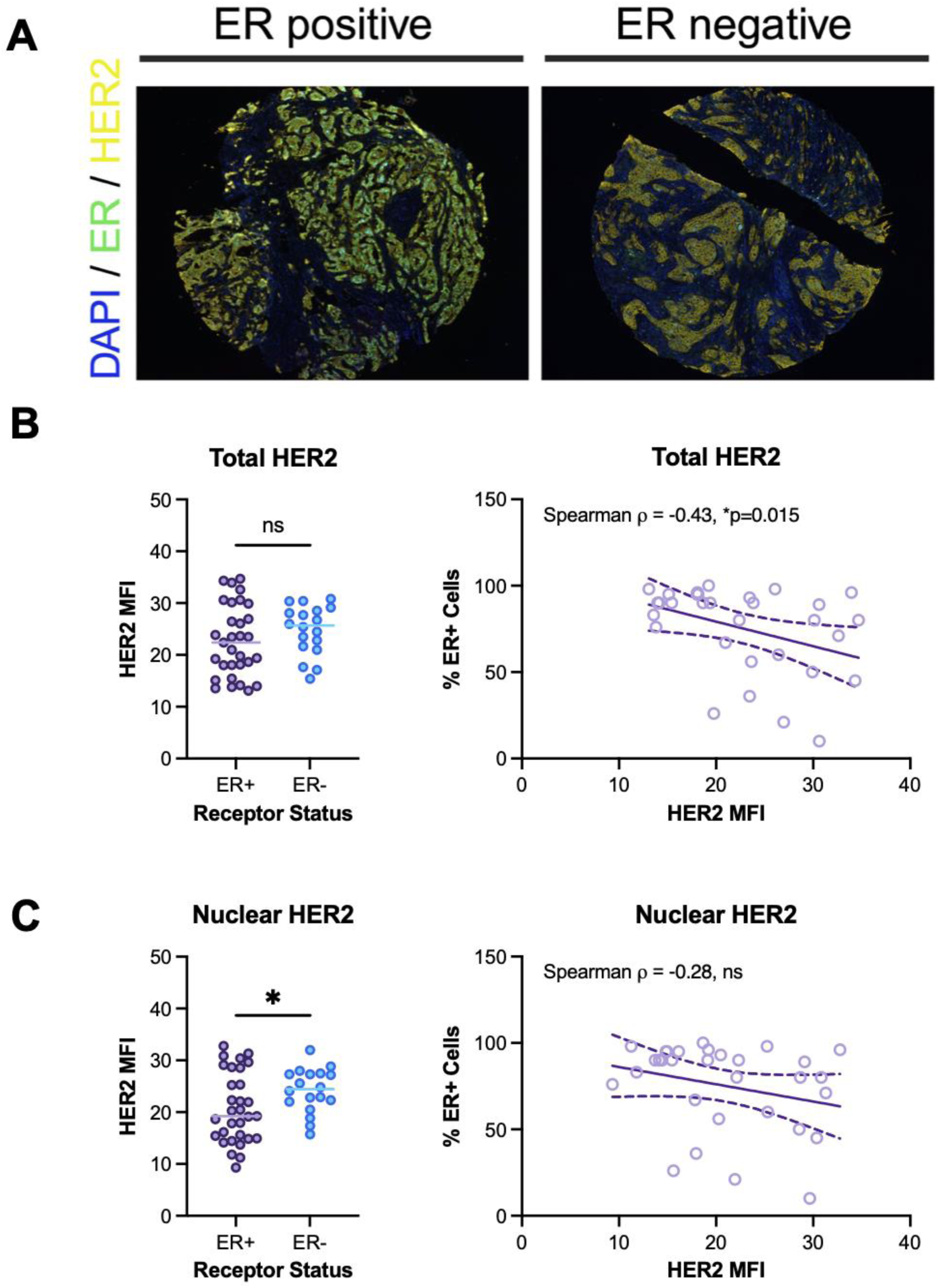
ER levels inversely correlate with HER2 in treatment-naive, HER2-amplified breast tumors. (A) Representative multiplex IHC staining of HER2 and ER for ER positive (ER+) and ER negative (ER-) breast tumor cores from a HER2+ treatment-naive breast cancer tissue microarray. HER2 signal is yellow, ER is green, DAPI/nuclei are blue. (B) Total or (C) nuclear HER2 mean fluorescence intensity (MFI) in pan-cytokeratin-positive tumor cells separated by tumor ER status (left panels), and Spearman correlation of percent ER+ cells with total/nuclear HER2 MFI (right panels). \**p* < 0.05; ns, not significant.

### Estrogen promotes semi-distinct ER-activated gene expression programs in ER+/HER2+ versus ER+ breast cancer cells

The negative correlation between ER and nuclear HER2 prompted us to investigate the role of ER in gene regulation in ER+/HER2+ breast cancer. While ER transcriptional regulatory programs have been well-defined in ER+ breast cancer, they have not been comprehensively assessed in ER+/HER2+ breast cancer. Therefore, we performed ER ChIP-seq to define the ER cistrome in the ER+/HER2+ breast cancer cell line BT-474. Hormone deprivation followed by 17-beta estradiol (E2) stimulation induced a robust, 3-fold increase in the number and more localized enrichment of the ER ChIP-seq peaks (Fig. 2A-2B). ChIP-qPCR further validated significant E2-induced ER recruitment to the promoter of the classical ER target gene *TFF1* in BT-474 and ER+ MCF-7 breast cancer cells (Fig. 2C).

**Figure 2.**
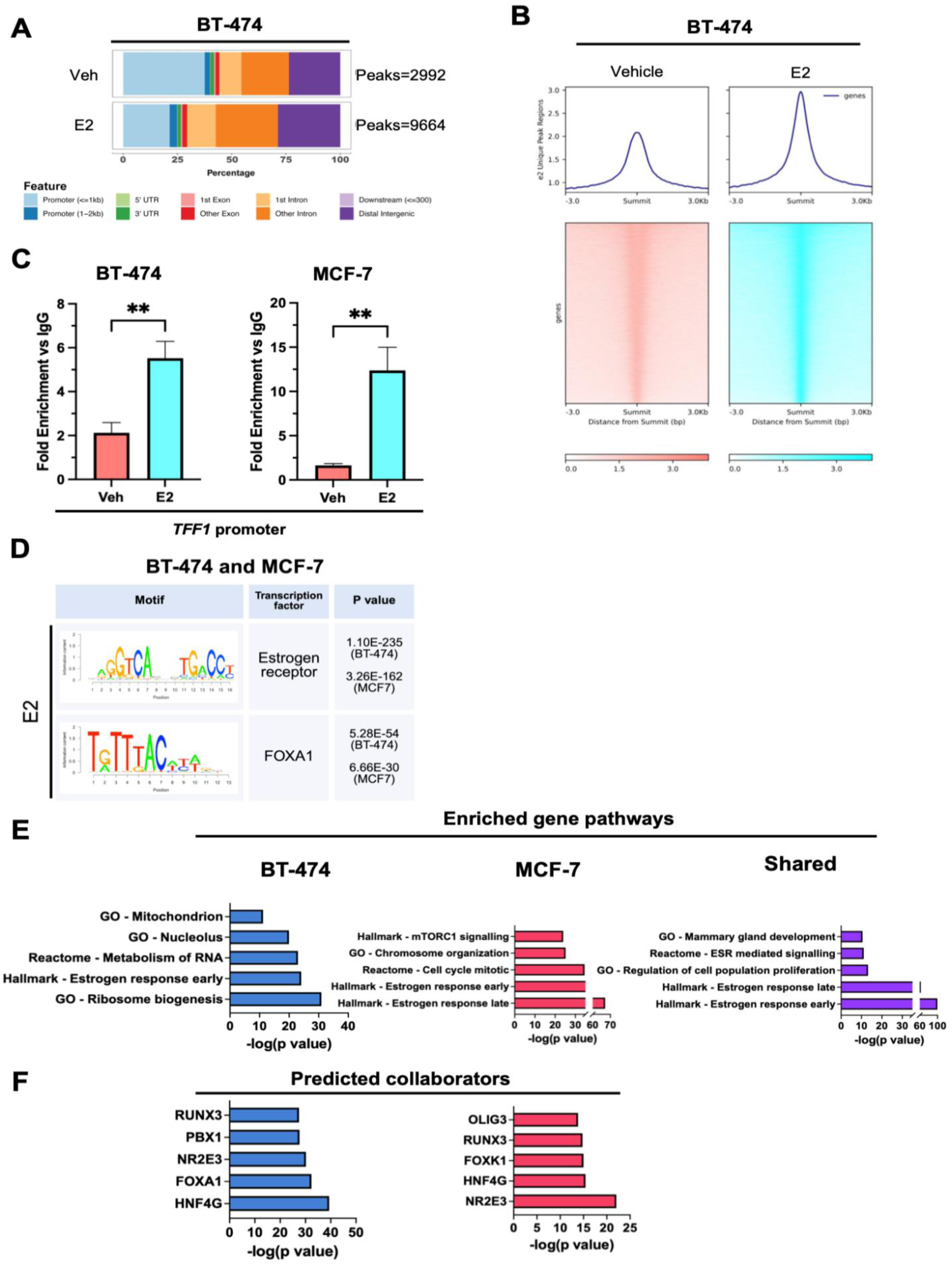
Estrogen activates common and specific transcriptional programs in ER+ and ER+/HER2+ breast cancer cells. (A) Genome binding locations and counts of ER ChIP-seq peaks in ER+/HER2+ BT-474 cells that were estrogen deprived for 3 days and stimulated with 10 nM E2 for 3 hours. (B) Heatmap of BT-474 ER ChIP-seq signal at E2-induced peaks. (C) ChIP-qPCR of ER at the *TFF1* promoter in ER+/HER2+ BT-474 and ER+ MCF-7 cells estrogen deprived and stimulated with 10 nM E2 for 3 hours. Data from 1 representative biological replicate is presented as the mean of technical replicates ± SD and analyzed by unpaired t test. (D) E2-enriched DNA binding motifs identified by BETA integration of ER ChIP-seq and RNA-seq from BT-474 cells and previously published MCF-7 data from Guan et al. (E) GSEA of genes associated with E2 stimulation specific to BT-474, MCF-7, or shared between cell lines, and (F) predicted ER transcriptional collaborators using BETA. GO, gene ontology; Hallmark, hallmark gene sets. \*\**p* < 0.01.

To identify potential ER-regulated genes mediated by E2 stimulation, we performed Binding and Expression Target Analysis (BETA), integrating parallel ER ChIP-seq and RNA-seq data from our BT-474 cells and previously published data for MCF-7 from Guan et al. (28,29). The top enriched DNA binding motifs identified by BETA in both BT-474 and MCF-7 cells were estrogen response elements (EREs) for ER, and FOXA1 binding sites (Fig. 2D). ER was recruited to the *PGR*, *TFF1*, and *FOXC1* enhancers and the *TFF1* promoter after E2 stimulation in both BT-474 and MCF-7 cells (Fig. S1A). The top enriched transcriptional programs commonly shared between BT-474 and MCF-7 cells were related to estrogen and ER signaling (29) (Fig. 2E). BT-474 cells had an enrichment of genes related to ribosome biogenesis, RNA processing and a subset of estrogen-related genes, while MCF-7 cells had increased expression of cell cycle-related genes and a different subset of estrogen-related genes (Fig. 2E). BETA also analyzes DNA motifs near ER peaks adjacent to differentially expressed genes to identify potential ER collaborators. Several predicted collaborators were shared between the two estrogen-stimulated cell lines, including the transcription factors RUNX3 and HNF4G (Fig. 2F).

Though estrogen-stimulated gene expression programs were largely consistent between the two cell lines, BETA analysis also identified cell line-specific or preferential E2-induced ER binding and gene expression patterns, such as the RNA binding protein PUM2 in BT-474 cells and the cell cycle regulator cyclin D1 (CCND1) in MCF-7 cells. Estrogen deprivation followed by E2 stimulation increased ER binding near *CCND1* in MCF-7 cells (Fig. S1B-S1C). ER binding near *PUM2* was increased by E2 in both cell lines (Fig. S1B-S1C. However, *PUM2* was not considered an ER- regulated gene in MCF-7 cells by BETA due to a lack of increased RNA expression with E2 stimulation (data not shown). In summary, while E2-stimulated ER regulates the expression of some unique genes in ER+/HER2+ breast cancer cells, it induces estrogen-related gene expression programs broadly similar to those found in ER+ breast cancer cells in the absence of HER2 amplification.

### Trastuzumab and pertuzumab promote classical ER target gene expression in ER+/HER2+ breast cancer cells

After establishing the similarities in ER gene regulation between ER+/HER2+ and ER+ breast cancer cells, we next addressed how targeted therapy affects ER and HER2 localization and function. Specifically, we assessed the effect of anti-HER2 targeted therapy on ER protein expression, HER2 protein expression and tyrosine phosphorylation, and classical ER target gene expression in ER+/HER2+ BT-474 cells. The selective ER degrader fulvestrant (ICI), serving as a control, decreased ER levels in whole cell lysate as expected, with little-to-no impact on HER2 expression or phosphorylation (Fig. 3A). Neither the combination of anti-HER2 monoclonal antibodies trastuzumab and pertuzumab (TP) nor the HER2-selective tyrosine kinase inhibitor (TKI) tucatinib (Tuc) had any substantive impact on ER or HER2 protein expression after three hours of treatment (Fig. 3A).

**Figure 3.**
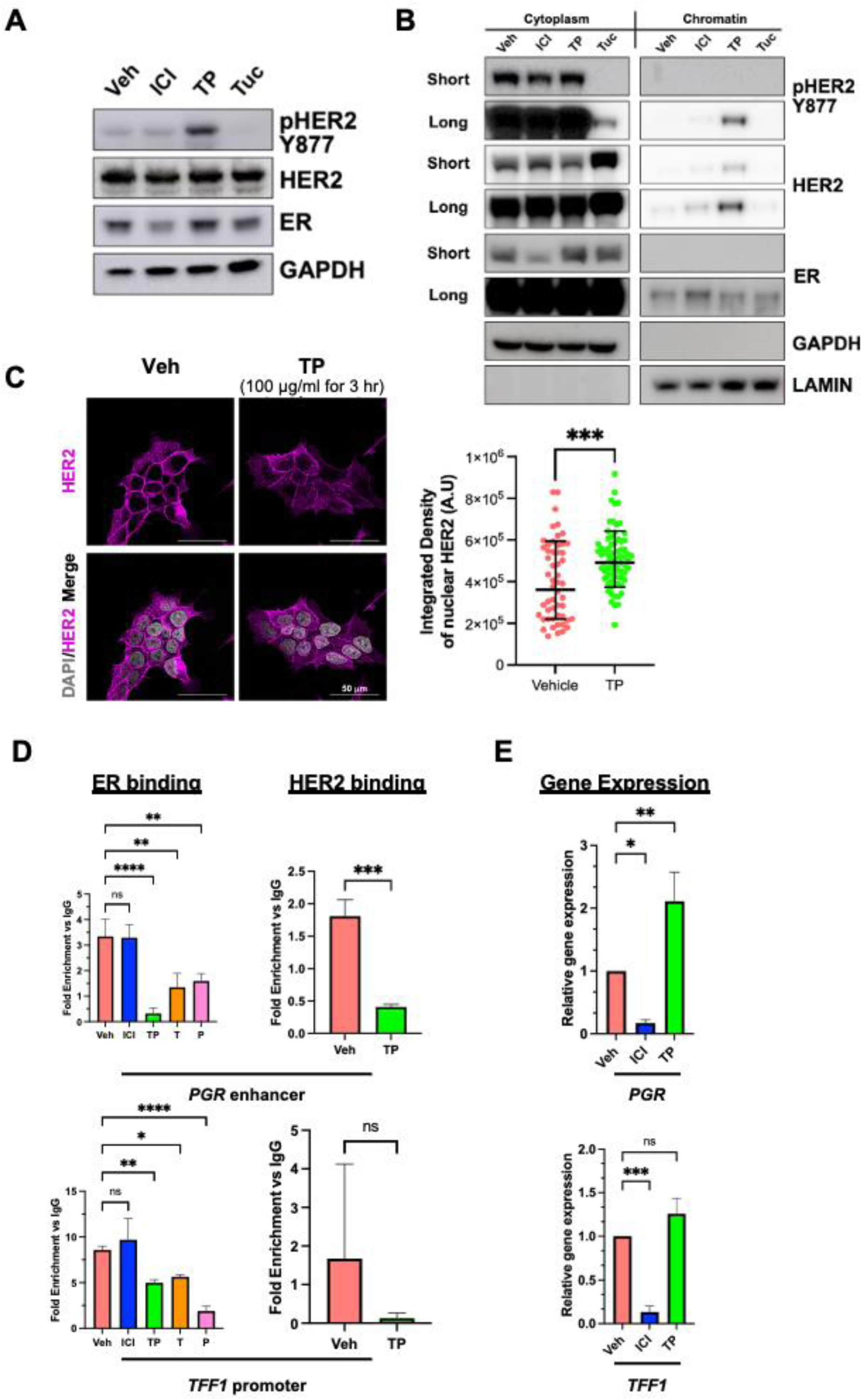
TP promotes expression of classical ER target genes in ER+/HER2+ breast cancer cells. (A) Immunoblot showing whole cell lysates from ER+/HER2+ BT-474 breast cancer cells treated with vehicle control (veh), 100 nM fulvestrant (ICI), 100 µg/ml trastuzumab and pertuzumab (TP), or 100 nM tucatinib (Tuc) for 3 hours. Phospho-HER2 (pHER2) Y877 denotes HER2 phosphorylation at tyrosine 877, GAPDH is the loading control. (B) Immunoblot of cytoplasmic and chromatin fractions of BT-474 cells treated as in (A). GAPDH and Lamin B1 are the loading controls for the cytoplasmic and chromatin-bound fractions, respectively. (C) Representative images of HER2 immunofluorescence staining and quantification of nuclear HER2 in BT-474 cells treated with TP for 3 hours from two independent biological replicates. HER2 is in magenta and DAPI is in gray. An unpaired t test was used. (D) ChIP-qPCR of ER or HER2 binding at the *PGR* enhancer and *TFF1* promoter in BT-474 cells treated with ICI, TP, 100 µg/ml trastuzumab (T), or 100 µg/ml pertuzumab (P). Data from 1 representative biological replicate is presented as a mean of technical replicates ± SD and analyzed by one-way ANOVA with Dunnett’s multiple comparisons test. (E) RNA expression of ER target genes *PGR* and *TFF1* of BT-474 cells treated for 24 hours with ICI or TP. Data from 3 independent biological replicates is presented as the mean ± SD and analyzed by one-way ANOVA with Dunnett’s multiple comparisons test. \**p* < 0.05, \*\**p* < 0.01, \*\*\**p* < 0.001, and \*\*\*\**p* < 0.0001; ns, not significant.

However, TP markedly increased phosphorylation of HER2 at two post-translational modification (PTM) sites that indicate an active receptor: tyrosine 877 within the kinase domain, and the tyrosine 1248 autophosphorylation site in the C-terminal cytoplasmic tail (Fig. 3A, S2A). In contrast, tucatinib reduced HER2 tyrosine 877 and tyrosine 1248 phosphorylation in whole cell lysates (Fig. 3A, S2A). Consistent with our observations, phosphorylation of HER2 at these residues has been reported with trastuzumab alone or the TP combination *in vitro* (50,51).

Next, we used cell fractionation to assess the effect of anti-HER2 targeted therapy on HER2 subcellular localization, since active HER2 dimers have been shown to translocate to the nucleus, and trastuzumab antibodies can also be internalized (52,53). The chromatin localization of tyrosine 877 pHER2 and total HER2 increased following treatment with TP, but not with tucatinib (Fig. 3B). Fulvestrant had no demonstrable effect on pHER2 or HER2 abundance in the cytoplasmic or chromatin fractions, but did decrease cytoplasmic ER concomitant with increased chromatin-associated ER, consistent with a prior study (29). Interestingly, trastuzumab or pertuzumab individually induced chromatin localization of tyrosine 877 pHER2 and total HER2 despite binding to different regions of HER2 (54), but this was not observed following treatment with epidermal growth factor (EGF, Fig. S2B). Unlike the TP combination, individual anti-HER2 antibodies did not promote chromatin localization of tyrosine 1248 pHER2 (Fig. S2C). We observed similar TP-induced tyrosine 877 pHER2 and total HER2 chromatin localization in the ER-, HER2 amplified cell line SK-BR-3, but not in ER+, non-HER2 amplified MCF-7 breast cancer cells (Fig. S2D), suggesting that TP-mediated chromatin localization of tyrosine 877 pHER2 and total HER2 is not dependent on the presence of ER. Finally, to corroborate these biochemical studies, we performed immunofluorescence in BT-474 cells to visualize HER2 nuclear localization in response to TP treatment, and found that TP treatment significantly increased nuclear HER2 levels (Fig. 3C). In summary, these data demonstrate that, alone or in combination, the anti-HER2 therapeutic antibodies TP, but not tucatinib, increased tyrosine 877 phosphorylation and chromatin localization of HER2.

Given the established crosstalk between HER2 and ER and our observations of TP induction of HER2 nuclear and chromatin localization, we assessed if TP affects ER chromatin binding at classical ER target genes using ChIP-qPCR. Acute treatment (three hours) with TP, or either trastuzumab or pertuzumab alone, significantly decreased ER occupancy at the *PGR* enhancer and *TFF1* promoter in BT-474 cells (Fig. 3D, left panels). Fulvestrant did not reduce ER occupancy at these loci, though it did reduce ER binding at these sites in ER+ MCF-7 cells (Fig. S2E). Because we observed increased HER2 chromatin localization with TP treatment, and prior studies have reported that HER2 is a chromatin-associated factor whose cistrome is expanded in response to activation by EGF (55), we also performed ChIP-qPCR for HER2 at these sites. Despite greater nuclear/chromatin localization of HER2, TP treatment significantly reduced HER2 binding at the *PGR* enhancer (Fig. 3D, top right panel). TP had no significant impact on HER2 binding at the *TFF1* promoter in BT-474 cells (Fig. 3D, bottom right panel), or either locus in ER-, HER2 amplified SK-BR-3 cells (Fig. S2E). It is important to note, however, that HER2 binding at these sites was minimal or absent (i.e. not enriched versus the IgG control) under vehicle-treated conditions.

We then determined if reduced ER binding was associated with altered mRNA expression of the classical ER target genes in TP-treated ER+/HER2+ BT-474 cells. TP significantly increased *PGR* expression, while fulvestrant caused a decrease in both *PGR* and *TFF1* expression as expected (Fig. 3E). *PUM2* expression, while not inhibited by fulvestrant, was significantly increased by TP treatment (Fig. S2F). Taken together, TP promotes classical ER target gene expression in ER+/HER2+ breast cancer cells, despite an acute reduction of ER genomic binding at associated enhancer and promoter sites.

### Resistance to trastuzumab and pertuzumab is associated with reduced ER binding to classical ER target genes

Long-term treatment with trastuzumab and the subsequent acquisition of resistance are associated with global transcriptomic alterations in HER2+ breast cancer cells (56,57). Therefore, we sought to characterize how resistance to TP may impact ER and HER2 subcellular localization and chromatin binding, as well as classical ER target gene expression, in ER+/HER2+ breast cancer cells. MDA-MB-361 is an ER+/HER2+ breast cancer cell line that is inherently (*de novo*) resistant to anti-HER2 monoclonal antibodies and tyrosine kinase inhibitors (58,59)(Fig. S3A). To model acquired TP resistance, we continuously cultured BT-474 cells in low dose TP for ∼3 months to generate a variant that is resistant to TP, but not cross-resistant to tucatinib (Fig. 4A). Acquired TP resistance was associated with a non-statistically significant increase in ER RNA (*ESR1*) and significant decrease in HER2 RNA (*ERBB2*, Fig. S3B). MDA-MB-361 cells also showed significantly elevated *ESR1* and reduced *ERBB2* expression compared to BT-474 cells (Fig. S3B) consistent with our prior report at the protein level (11).

**Figure 4.**
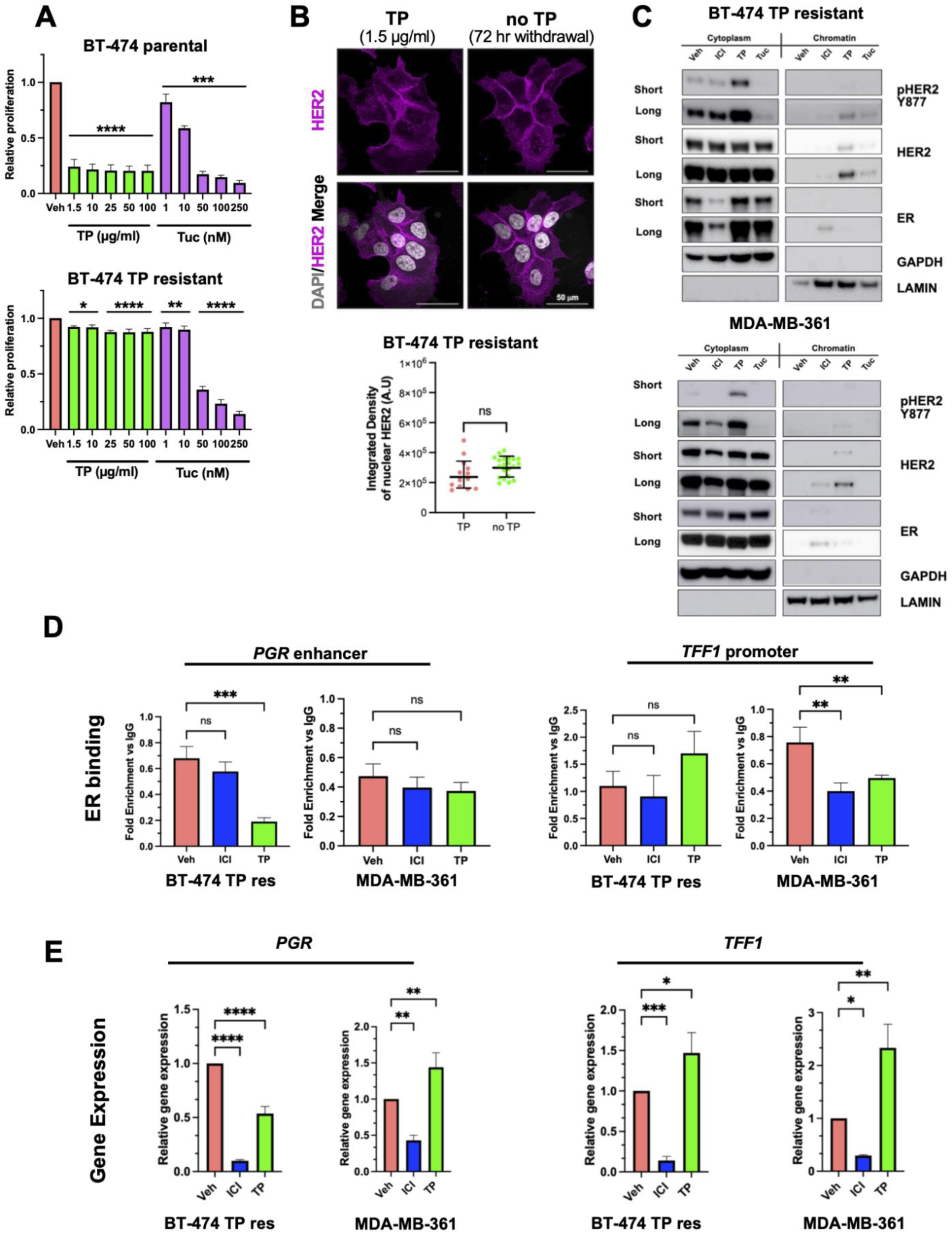
Resistance to TP is associated with reduced ER binding to classical ER target genes relative to sensitive counterparts. (A) Crystal violet assays of BT-474 and BT-474 TP resistant cells treated with the indicated concentrations of TP or Tuc for 8 days. One-way ANOVA with Dunnett’s multiple comparisons test was used to compare each treatment concentration with vehicle (Veh) control; asterisks denote significance and are defined below. (B) Representative images of HER2 immunofluorescence staining and quantification of nuclear HER2 in BT-474 TP resistant cells maintained in 1.5 µg/ml TP media or after a 72-hour TP washout from 2 biological replicates; unpaired t test was used. (C) Immunoblots of cytoplasmic and chromatin fractionations of BT-474 TP resistant cells and TP-resistant ER+/HER2+ MDA-MB-361 breast cancer cells, treated with 100 nM ICI, 100 µg/ml TP, or 100 nM Tuc for 3 hours. (D) ChIP-qPCR of ER binding at the *PGR* enhancer and *TFF1* promoter in BT-474 TP resistant cells and MDA-MB-361 cells. Data from 1 representative biological replicate is presented as the mean of technical replicates ± SD and analyzed by unpaired t test. (E) RNA expression of *PGR* and *TFF1* in BT-474 TP resistant and MDA-MB-361 cells treated for 24 hours. Data from 3 independent biological replicates is presented as the mean ± SD and analyzed by one-way ANOVA with Dunnett’s multiple comparisons test. \**p* < 0.05, \*\**p* < 0.01, \*\*\**p* < 0.001, and \*\*\*\**p* < 0.0001; ns, not significant.

In BT-474 TP resistant cells, nuclear HER2 localization was observed by immunofluorescence, and withdrawal of TP for 3 days did not alter the levels of nuclear HER2 (Fig. 4B). Cell fractionation analysis showed that TP moderately increased HER2 and tyrosine 877 pHER2 chromatin localization in both BT-474 TP resistant and MDA-MB-361 cells (Fig. 4C). Similar to BT-474 parental cells, tucatinib did not induce HER2 chromatin localization, and fulvestrant did not impact pHER2 or HER2 in the cytoplasmic or chromatin fractions, but increased chromatin-associated ER in both resistant cell lines.

In contrast to BT-474 parental cells that demonstrated robust ER recruitment to the *PGR* enhancer and *TFF1* promoter at baseline that was significantly attenuated by TP treatment (Fig. 3D), both TP-resistant cell lines showed little or no ER binding at these sites under vehicle-treated conditions (fold enrichment versus the IgG control <1, Fig. 4D). Similar to BT-474 parental cells, HER2 exhibited minimal or no binding to the *PGR* enhancer or *TFF1* promoter in BT-474 TP resistant cells (Fig. S3C). At the RNA level following 24 hours of treatment, BT-474 TP-resistant cells no longer exhibited TP-induced *PGR* expression but showed TP-mediated induction of *TFF1*, whereas MDA-MB-361 cells displayed significant TP-induced expression of both *PGR* and *TFF1* (Fig. 4E). Together, these data suggest that HER2 and pHER2 chromatin accumulation, and induction of classical ER target gene expression, can still occur in response to TP in ER+/HER2+ breast cancer cells with either acquired or *de novo* resistance. However, ER binding to selected classical target gene promoters and enhancers at baseline is notably reduced in both TP-resistant models compared to TP-responsive cells.

### Trastuzumab and pertuzumab promote ER binding to sites near genes associated with worse outcomes in HER2+ breast cancer

The reduction of ER binding at ER target genes in TP-resistant cells led us to profile the global ER cistrome in response to anti-HER2 therapy. We performed CUT&RUN assays to determine whether TP can elicit genome-wide alterations in the ER cistrome in BT-474 cells. Acute (3 hour) treatment with TP dramatically increased the number of ER peaks across the genome, and shifted the top enriched ER-bound motifs from the estrogen and other nuclear receptor response elements toward motifs corresponding to zinc finger transcription factors (Fig. 5A-5B). Pathway analysis of genes assigned to ER peaks unique to the TP-treated condition showed significant enrichment of several signaling pathways, including the Wnt and RAGE pathways (Fig. 5C). Genes within these pathways included *FZD8*, which encodes the Frizzled-8 receptor of the Wnt pathway, and *RELA*, which encodes the p65 subunit of NF-kB and is an effector of the RAGE pathway (60,61)(Fig. 5D). Of note, TP treatment led to ER recruitment near two established E2 inducible genes, *CCND1* and *BCL2* (62,63). However, the sequence motifs underlying these TP-induced peaks corresponded to zinc finger transcription factors, not EREs, suggesting that ER binding at these genes may be mediated by tethering to other transcription factors.

**Figure 5.**
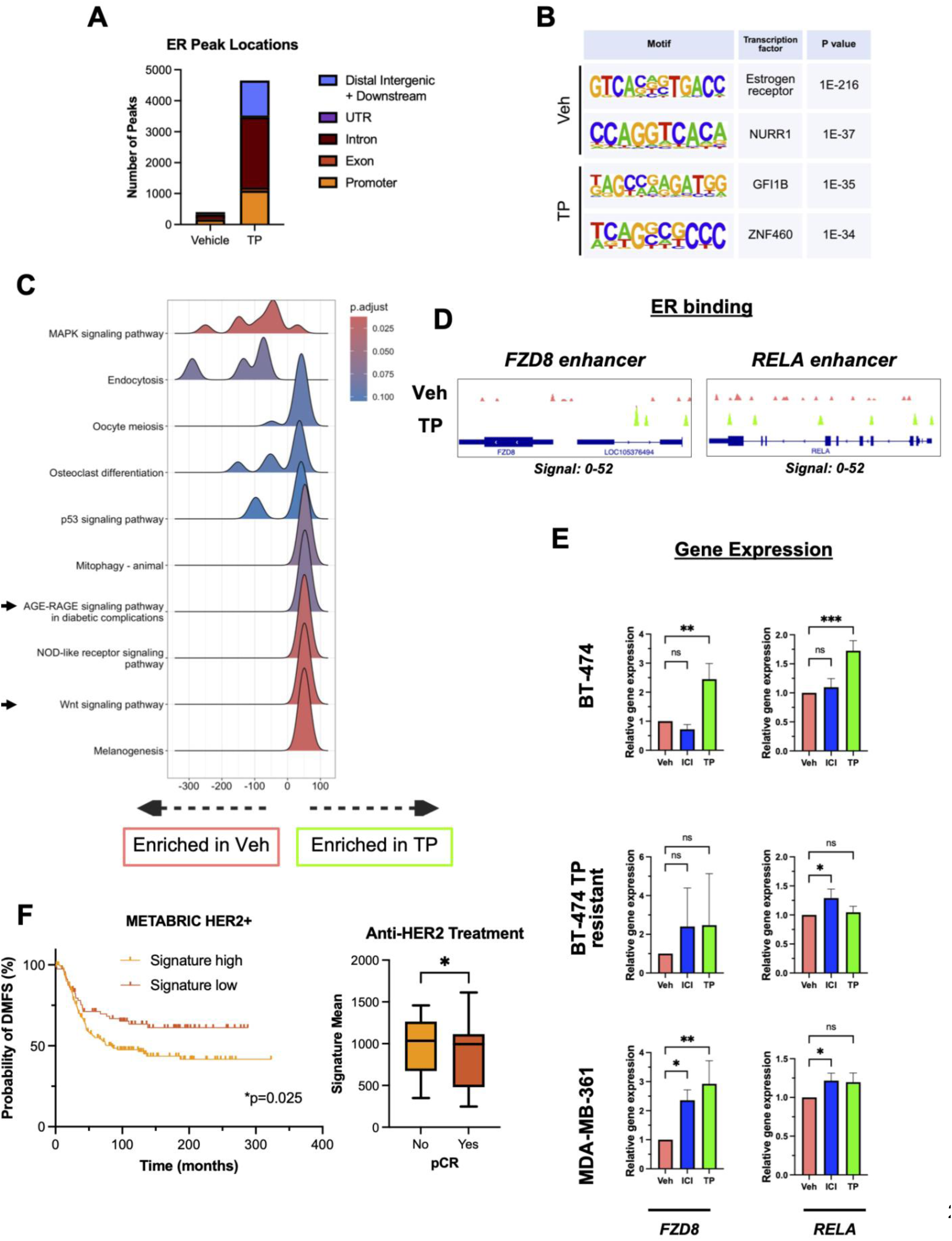
TP induces gain of ER binding to novel target genes, and their elevated expression correlates with worse DMFS and failure to achieve pCR. (A) Genome binding locations and counts of ER CUT&RUN peaks in BT-474 cells treated with TP for 3 hours. (B) DNA binding motifs associated with ER-bound sites in BT-474 cells identified by HOMER de novo motif analysis. (C) Gene set enrichment pathway analysis of genes assigned to ER peaks in vehicle and TP-treated conditions of BT-474 cells. Gene sets with a peak score above 0 are enriched in TP-treated cells, whereas gene sets below 0 are enriched in vehicle-treated cells. (D) ER binding at the *FZD8* (Wnt pathway) and *RELA* (AGE-RAGE pathway) genes. (E) RNA expression of *FZD8* and *RELA* in BT-474, BT-474 TP resistant, and MDA-MB-361 cells treated with ICI or TP for 24 hours. Data from 3 independent biological replicates is presented as the mean ± SD and analyzed by one-way ANOVA with Dunnett’s multiple comparisons test. (F) Distant metastasis-free survival (DMFS) curves of 234 patients with HER2+ breast cancer in the METABRIC cohort, separated by high (orange, n=153) and low (red, n=81) median expression of the 39-gene ER-bound signature derived from the top 5 enriched pathways in (C; genes listed in Table S4); a log-rank Mantel-Cox test was used. In a separate cohort of 123 patients with HER2+ breast cancer treated with anti-HER2 therapy (46), expression of the same signature was compared between patients with (yes, n=56) and without (no, n=67) pathological complete response (pCR) using Mann-Whitney test. \**p* < 0.05, \*\**p* < 0.01, and \*\*\**p* < 0.001; ns, not significant.

Assessment of *FZD8*, *RELA*, *BCL2*, and *CCND1* mRNA levels by qPCR following longer term (24 hour) treatment with TP showed significantly increased expression of all four genes in parental BT-474 cells, and all except *RELA* in *de novo* TP-resistant MDA-MB-361 cells (Fig. 5E, S4A).

However, none of the four genes were induced by TP in the acquired TP resistance BT-474 model, and fulvestrant did not suppress expression of any of the four genes in any cell line. Together these data demonstrate that despite promoting loss of ER binding at classical ER target genes *TFF1* and *PGR*, TP broadly reprograms the ER cistrome, increasing its genomic binding at zinc finger motifs and genes associated with Wnt and RAGE signaling. Additionally, TP treatment significantly increases mRNA expression of these novel targets in TP-responsive and resistant ER+/HER2+ cell lines.

To determine whether genes associated with TP-induced ER peaks were linked to clinical outcome measures, we first established a signature of 39 unique genes from the top 5 enriched pathways (Fig. 5C, Table S4). We then determined the relationship of this signature (and select individual genes of interest) to survival and pCR measures in multiple clinical datasets. High expression (above median) of the 39-gene signature was significantly associated with reduced distant metastasis-free survival (DMFS, Fig. 5F, left panel) in 234 patients with HER2+ breast cancer from the METABRIC database. This relationship between signature expression and DMFS was not observed in ER+ breast cancer (Fig. S4B). We then queried publicly available matched transcriptomic and treatment response data for a pooled dataset of 123 HER2+ breast cancers from patients treated with anti-HER2 therapy (46). Mean expression of the 39-gene signature was significantly elevated in those individuals who did not achieve pCR following neoadjuvant anti-HER2 therapy (Fig. 5F, right panel). Individual expression of *FZD8* or *CCND1* was also significantly increased in those who did not achieve pCR (data not shown).

Lastly, we determined whether expression of these TP-induced putative ER target genes was associated with pCR in the RNA-seq cohort from NSABP B-41 (n=245; ER+ = 141, ER- = 104), a Phase 3 randomized clinical trial that evaluated neoadjuvant chemotherapy combined with HER2- targeted therapies in patients with HER2+ breast cancer, with pCR as the primary outcome (14). Prior correlative studies in B-41 have established that individual *ERBB2* and *ESR1* expression, and several immune-related signatures, are predictive of pCR status (47,64). In B-41, we found increased *FZD8* or *BCL2* expression in tumors of patients who did not achieve pCR in the overall cohort (Fig. S4C).

Altogether, these findings demonstrate that TP induces ER binding to novel target genes in ER+/HER2+ breast cancer cells, and that elevated expression of a subset of these genes correlates with worse DMFS in METABRIC, and residual disease following neoadjuvant anti-HER2 therapy in B- 41, in HER2+ breast cancers.

## Discussion

Although anti-HER2 monoclonal antibody therapy with TP is highly effective, co-expression of ER or another HR reduces pCR rates (13–18). Here, we demonstrate that HER2 inhibition by TP can actively reprogram ER, which may ultimately contribute to adaptive resistance in ER+/HER2+ breast cancer. In treatment-naive tumors, nuclear HER2 inversely correlates with ER, while *in vitro*, acute TP treatment promotes HER2 chromatin accumulation while simultaneously displacing ER from canonical targets. Genome-wide profiling revealed that TP triggers a profound redistribution of ER binding toward non-canonical zinc finger motifs, adjacent to genes in the Wnt and RAGE pro-survival signaling pathways. Elevated expression of this gene signature strongly correlates with shorter DMFS and reduced likelihood of achieving pCR in HER2+ breast cancer clinical cohorts. These collective findings provide a mechanistic rationale highlighting the importance of targeting both ER and HER2 simultaneously alongside additional pro-survival pathways, and support the recent FDA approval of CDK4/6 inhibitor palbociclib in combination with trastuzumab (with or without pertuzumab) and endocrine therapy as maintenance therapy for ER+/HER2+ locally advanced or metastatic breast cancer, following the successful PATINA trial (65).

Our analysis of a cohort of treatment-naive ER+/HER2+ primary tumors identified an inverse correlation between nuclear HER2 expression and ER levels, with higher nuclear HER2 in ER- compared to ER+ tumors. This inverse relationship between HR and nuclear HER2 may align with recent molecular subtyping defining distinct HER2-dominant (MUKDEN I) versus ER-dominant (MUKDEN II) subgroups in ER+/HER2+ disease (66). Nuclear HER2 has previously been linked to shorter DMFS in tamoxifen-treated ER+ (non-HER2-amplified) breast cancer (67), and nuclear HER2- EGFR dimers can promote progression of ductal carcinoma *in situ* toward a more malignant phenotype (52). Further studies in larger, clinically annotated ER+/HER2+ patient cohorts will be required to assess the impact of nuclear HER2 expression on response to targeted therapies.

We observed nuclear and chromatin-associated HER2 to be present in TP-treated ER+/HER2+ BT-474 cells and ER-/HER2+ SK-BR-3 cells, but not in non-HER2-amplified MCF-7 cells, suggesting that endogenously high levels of HER2 expression are required for this phenotype. Additionally, TP (but not tucatinib or EGF) specifically induced chromatin accumulation of total and Y877-phosphorylated HER2. This is consistent with prior studies, in which TP have been reported to promote HER2 phosphorylation *in vitro* (50,51). Our finding that tucatinib fails to induce HER2 movement suggests that HER2 kinase activity and Y877 phosphorylation facilitate this nuclear shift. The absence of pY1248 HER2 in the chromatin fraction in our studies may further suggest a selective role for Y877 over Y1248 in governing nuclear localization. However, site-directed mutagenesis targeting Y877 and Y1248 will be necessary to confirm whether Y877 phosphorylation is truly required for HER2 nuclear trafficking and chromatin association.

Prior studies have established that nuclear HER2 can act as a direct transcriptional coregulator at specific promoters, such as COX-2 (encoded by *PTGS2*) and ribosomal RNA genes (68,69), or across EGF-regulated genomic sites (55). However, despite TP-induced movement of HER2 to chromatin, our ChIP-qPCR assays revealed minimal direct HER2 enrichment at classical ER target loci like *PGR* and *TFF1*, suggesting that nuclear HER2 does not broadly co-occupy canonical ER target sites. Future studies utilizing proximity ligation assays, mass spectrometry, or genome-wide HER2 profiling will be needed to determine whether TP-stimulated chromatin-bound HER2 associates with other transcriptional complexes (such as those involving STAT3) or physically interacts with ER in the nuclear or chromatin compartment.

Acute (3-hour) TP treatment significantly reduced ER genomic occupancy at classical target gene loci (*PGR* and *TFF1*), whereas extended 24-hour TP treatment paradoxically increased *PGR* RNA expression. Of note, *PGR* mRNA was not increased by 6- or 9-hour TP treatment. This temporal decoupling, whereby acute ER displacement from canonical EREs is followed by later target gene upregulation, suggests that drivers of later ER target gene expression at 24 hours are independent of the near-term loss of ER binding at their promoters and enhancers. Compensatory transcription factors binding at the same or different loci (induced by TP), or ER-bound or tethered to other regulatory sites of the same genes could further explain our observations. Time-course cistromic and transcriptomic analyses across intermediate timepoints will be needed to elucidate the exact sequence of co-regulator assembly and chromatin remodeling that enables target gene induction following initial ER displacement.

Acquired TP resistance in BT-474 cells is characterized by HER2 chromatin localization in response to TP. This connects with established literature showing that HER2 endocytosis, recycling, and altered chromatin accessibility contribute to anti-HER2 antibody resistance (70–73). However, *in vitro* models of resistance cannot fully capture the immune-mediated mechanisms of anti-HER2 antibodies, such as antibody-dependent cellular cytotoxicity (ADCC) (74). *In vivo* modeling in immunocompetent systems will be essential to determine how tumor microenvironment (TME) factors and immune effector responses influence TP-induced HER2 chromatin localization and the ER cistrome in intact tumors. In our models of acquired (BT-474 TP resistant) and innate (MDA-MB-361) TP resistance, cells also exhibited reduced baseline ER and HER2 binding at classical ER target gene loci compared to sensitive BT-474 parental cells, yet retained TP-induced Y877 HER2 phosphorylation. The retained sensitivity to tucatinib in our acquired resistance model aligns with clinical evidence that HER2 kinase activity remains an actionable driver even after TP failure, such as the HER2CLIMB-05 trial demonstrating improved progression-free survival (PFS) with the addition of tucatinib (75).

Surprisingly, genome-wide ER CUT&RUN profiling demonstrated that acute TP treatment triggers a massive redistribution of the ER cistrome toward non-canonical zinc finger motifs adjacent to pro-survival genes (such as *FZD8*, *RELA*, *BCL2*, and *CCND1*), despite reducing ER occupancy at well-established promoter and enhancer loci. Importantly, longer-term TP treatment (24 hours) significantly induces mRNA expression of several of these genes, most notably *FZD8*. Additionally, the TP-induced ER cistrome appears to be distinct from growth factor-stimulated ER binding, whereby growth factor stimulation (such as EGF) has been shown to induce a distinct ER cistrome enriched for AP-1 motifs (76). Differences in enriched motifs between TP-induced ER target genes and EGF-stimulated ER targets suggest that anti-HER2 therapy reprograms ER via a mechanism distinct from classical growth factor crosstalk. Further study will be necessary to identify the specific zinc finger transcription factors that collaborate with ER to direct this TP-induced cistromic shift.

Expression of a 39-gene signature comprising the top TP-induced ER peaks was associated with shorter distant metastasis-free survival in METABRIC HER2+ breast cancers (Table S4). Higher expression of these genes was observed in tumors from patients not achieving pCR in response to anti-HER2 neoadjuvant therapy in a publicly available pooled cohort (46), with specific targets (*FZD8* and *BCL2*) also serving as pCR-predictive genes in NSABP B-41 (64). Of the 39 TP-induced genes, 4 correspond to the MUKDEN IV molecular subtype, which is characterized by pathway heterogeneity, Wnt signaling, and drug metabolism (66). Several of the genes we identify as exhibiting ER recruitment to regulatory sites (and increased mRNA expression) in response to TP are components of the RAGE signaling pathway. It has recently been reported that cancer-associated fibroblasts (CAFs) secreting S100-A11 can activate RAGE signaling to promote TP resistance (77), and TP can induce NF-κB signaling (78). TME-inclusive immunocompetent *in vivo* models and prospective clinical studies will be needed to evaluate whether targeting these TP-induced pathways (such as Wnt or RAGE/NF-κB signaling) can overcome or prevent resistance to anti-HER2 therapy in ER+/HER2+ breast cancer.

Our study has some limitations that may affect the interpretation of these findings. First, experiments were conducted in full serum (except for E2 stimulation experiments).In particular, although EGF has been shown to reduce the inhibitory effect of trastuzumab on BT-474 colony formation (79), EGF stimulation in our hands did not promote HER2 phosphorylation like the individual monoclonal antibodies or the TP combination did (Fig. S2B). Second, we did not fully explore the precise molecular machinery governing HER2 chromatin localization. While our results suggest that phosphorylation at the Y877 residue correlates with HER2 chromatin localization, whereas Y1248 phosphorylation does not, it remains unclear whether Y877 is the sole determinant of HER2 recruitment to chromatin. Previous studies have established that HER2 membrane trafficking and endosomal recycling regulate receptor concentration at the plasma membrane and modulate interactions with other ERBB family members, such as EGFR (80–82). Moving forward, integrating cell-surface trafficking dynamics with paired nuclear HER2 and ER cistromic profiling will be essential to fully map the transcriptional complexes triggered by anti-HER2 targeted therapies. Ultimately, defining how ER and HER2 dynamically co-regulate pro-survival transcription under therapeutic pressure will provide critical insights for designing novel combination strategies—such as optimized co-targeting or sequence-specific regimens combining anti-HER2 agents with next-generation endocrine therapies—to prevent or overcome treatment resistance in ER+/HER2+ breast cancer.

## Supplemental materials and methods

**Table S1.** List of antibodies.

| Name | Company | Catalog number | RRID | Usage |
| --- | --- | --- | --- | --- |
| phospho-HER2 Y877 | Cell Signaling | 2241 | AB 2099407 | Western blot |
| phospho-HER2 Y1248 | Cell Signaling | 2247 | AB 331725 | Western blot |
| HER2 | Cell Signaling | 2242 | AB 331015 | Western blot, ChIP |
| Estrogen receptor alpha | Abcam | ab32063 | AB 732249 | Western blot, ChIP |
| GAPDH | Proteintech | 60004-1-Ig | AB 2107436 | Western blot |
| Lamin B1 | Cell Signaling | 13435 | AB 2737428 | Western blot |
| HER2 | Cell Signaling | 2165 | AB 10692490 | Immunofluorescence |
| anti-RabbitIgG secondary, Alexa Fluor™ 594 | Thermo | A-11037 | AB 2534095 | Immunofluorescence |
| Estrogen receptor alpha | Millipore | 06-935 | AB 310305 | ChIP-seq |
| IgG control | EpiCypher | 13-0042 | AB 2923178 | CUT&RUN |
| H3K4me3 control | EpiCypher | 13-0041 | AB 3076423 | CUT&RUN |
| H3K27me3 control | EpiCypher | 13-0055 | AB 3665059 | CUT&RUN |
| Estrogen receptor alpha | EpiCypher | 13-2012 | AB 3720839 | CUT&RUN |

**Table S2.**
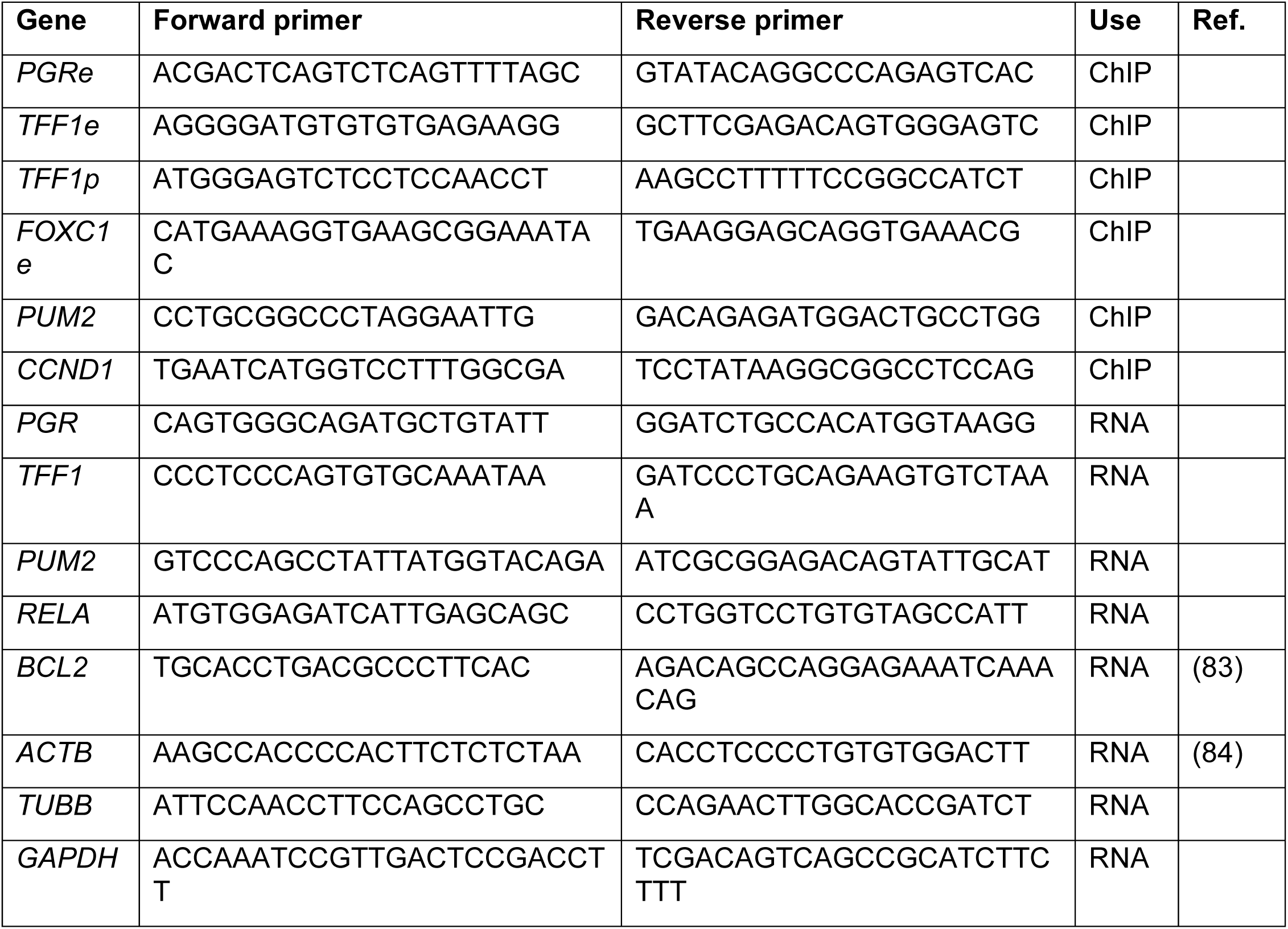
List of primer sequences.

**Table S3.**
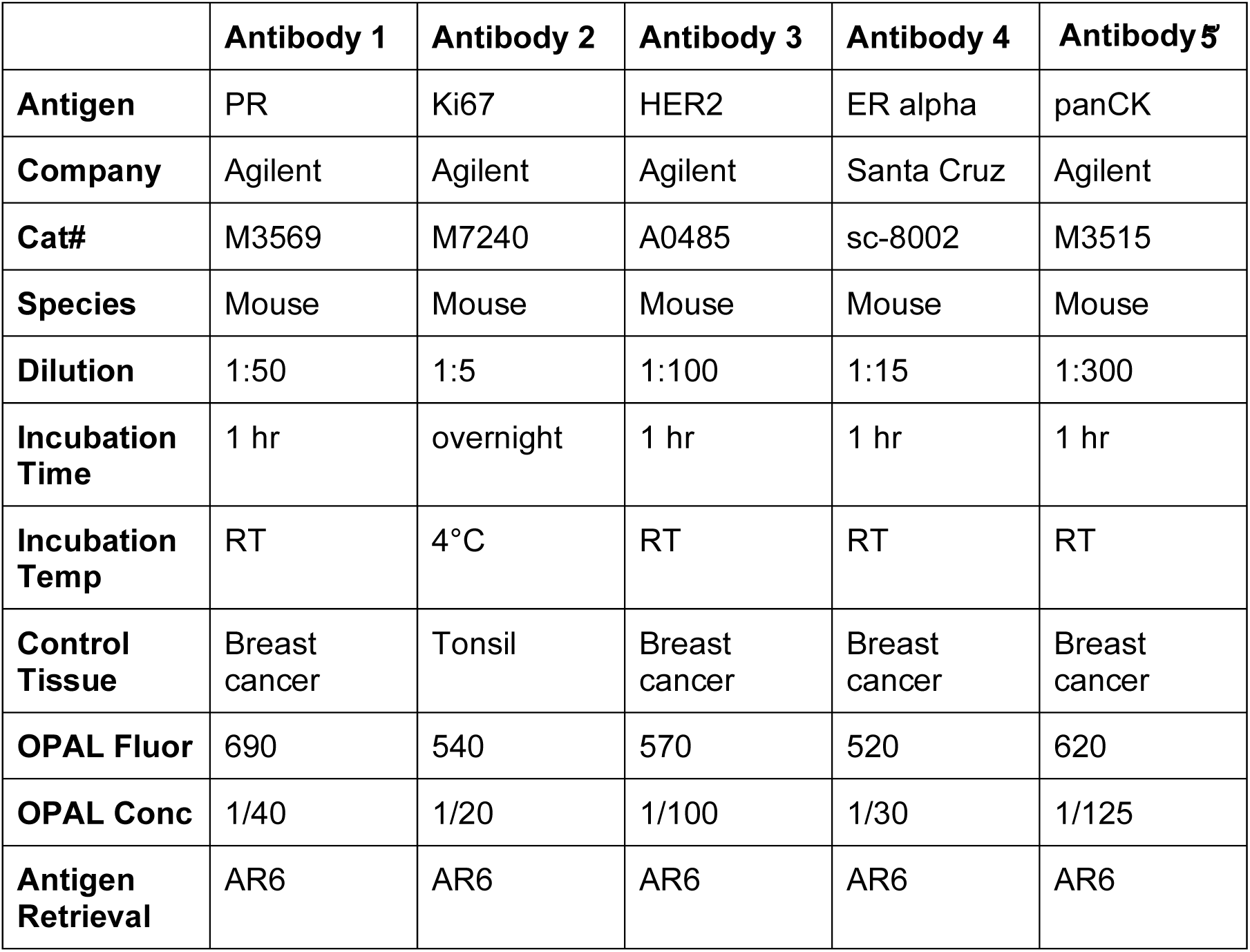
List of antibodies, OPAL dye pairings, and incubation conditions used for multiplex IHC of TMAs.

## Supplemental data

**Figure S1.**
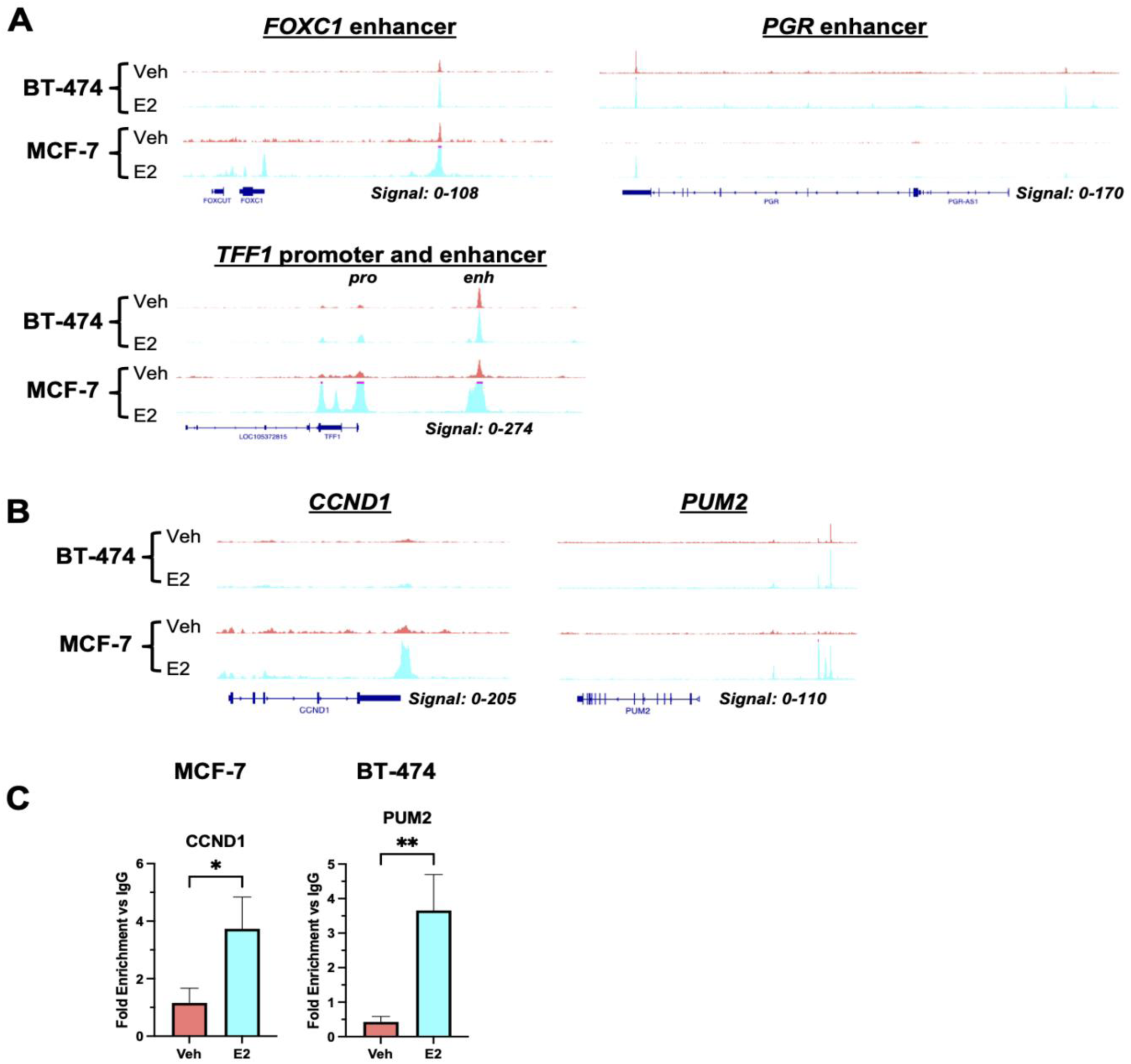
Estrogen receptor chromatin binding at target genes in ER+/HER2+ breast cancer cells. (A) ER ChIP-seq signal at the *FOXC1* enhancer, *PGR* enhancer, *TFF1* promoter, and *TFF1* enhancer of BT-474 and MCF-7 cells treated with E2 after estrogen deprivation. Areas in magenta represent signal that extends beyond the displayed numerical data range. Veh, vehicle control; E2, estradiol. (B) ER ChIP-seq signal at the *CCND1* and *PUM2* loci in BT-474 and MCF-7 cells identified by BETA. *CCND1* was considered an ER target gene in MCF-7, whereas *PUM2* was considered an ER target gene in BT-474 based on their significantly increased expression and changes in peak height. (C) ChIP-qPCR of ER at *CCND1* and *PUM2* loci in estrogen-deprived MCF-7 and BT-474 cells stimulated with 10 nM E2 for 3 hours. Data from 1 representative biological replicate is presented as the mean of technical replicates ± SD and analyzed by unpaired t test. pro = promoter, enh = enhancer, \**p* < 0.05 and \*\**p* < 0.01.

**Figure S2.**
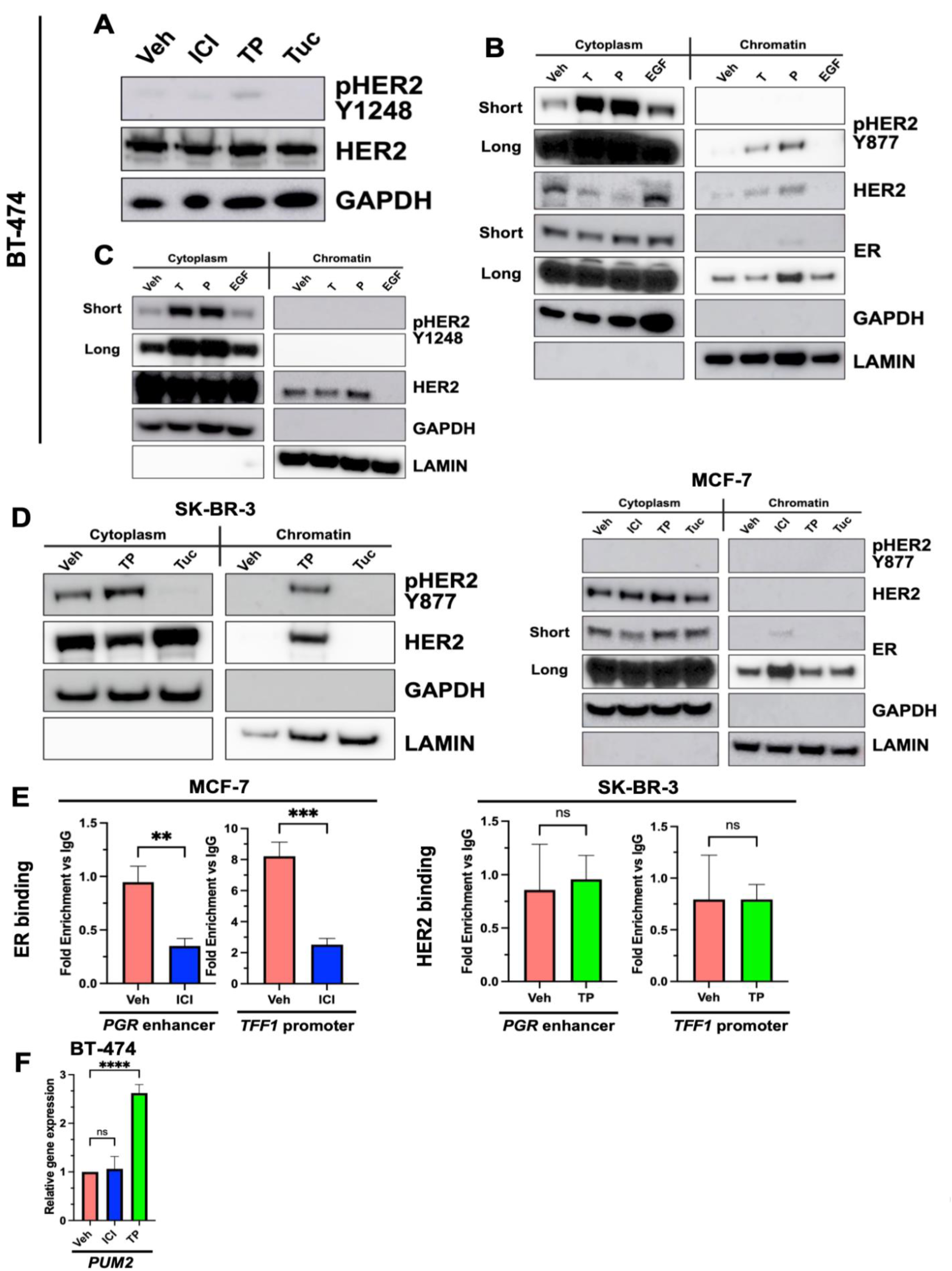
Effects of targeted therapies on ER and HER2 protein expression, localization, and chromatin binding in breast cancer cell lines. (A) Immunoblot showing whole cell lysates of BT-474 cells treated with 100 nM fulvestrant (ICI), 100 g/ml trastuzumab and pertuzumab (TP), or 100 nM tucatinib (Tuc) for 3 hours. pHER2 Y1248 denotes HER2 phosphorylation at tyrosine 1248. (B) and (C) Western blots of chromatin fractionation lysates from BT-474 cells treated with trastuzumab, pertuzumab, or EGF. (D) Chromatin fractionation immunoblot of HER2+ SK-BR-3 breast cancer cells treated with TP or Tuc, and ER+ MCF-7 breast cancer cells treated with ICI, TP or Tuc for 3 hours. (E) ChIP-qPCR of ER in MCF-7 cells, and HER2 in SK-BR-3, at the *PGR* enhancer and *TFF1* promoter. Cells were treated with ICI or TP for 3 hours. Unpaired t test was used. (F) RNA expression of *PUM2* in BT-474 cells treated with ICI or TP for 24 hours. One-way ANOVA with Dunnett’s multiple comparisons test was used. \*\**p* < 0.01, \*\*\**p* < 0.001, and \*\*\*\**p* < 0.0001; ns, not significant.

**Figure S3.**
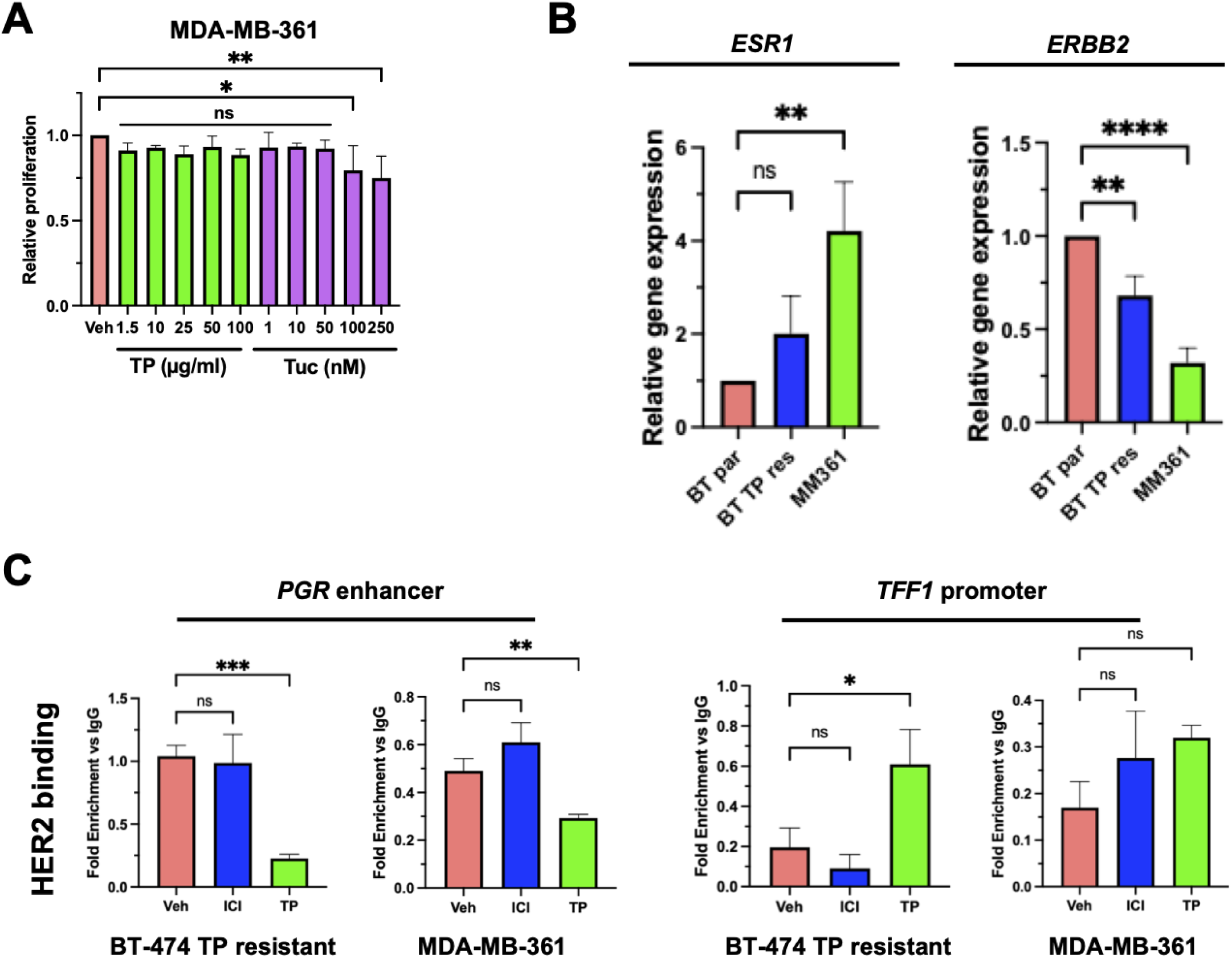
Characterization of HER2 expression and binding at classical ER target genes in intrinsic and acquired TP-resistant ER+/HER2+ breast cancer cells. (A) Crystal violet assays of TP-resistant MDA-MB-361 cells treated with the indicated concentrations of TP or Tuc for 8 days. One-way ANOVA with Dunnett’s multiple comparisons test was used to compare each treatment concentration to the vehicle (Veh) control. (B) RNA expression of *ESR1* and *ERBB2* of BT-474 parental (BT par), BT-474 TP-resistant (BT TP res), and MDA-MB-361 (MM361) cells. Data from 3 independent biological replicates is presented as the mean ± SD and analyzed by one-way ANOVA with Dunnett’s multiple comparisons test. (C) ChIP-qPCR of HER2 binding at the *PGR* enhancer and *TFF1* promoter in BT-474 TP-resistant and MDA-MB-361 cells following 3-hour treatments with ICI or TP. Data from 1 representative biological replicate is presented as the mean of technical replicates ± SD and analyzed by unpaired t test. \**p* < 0.05, \*\**p* < 0.01, \*\*\**p* < 0.001, \*\*\*\**p* < 0.0001; ns, not significant.

**Figure S4.**
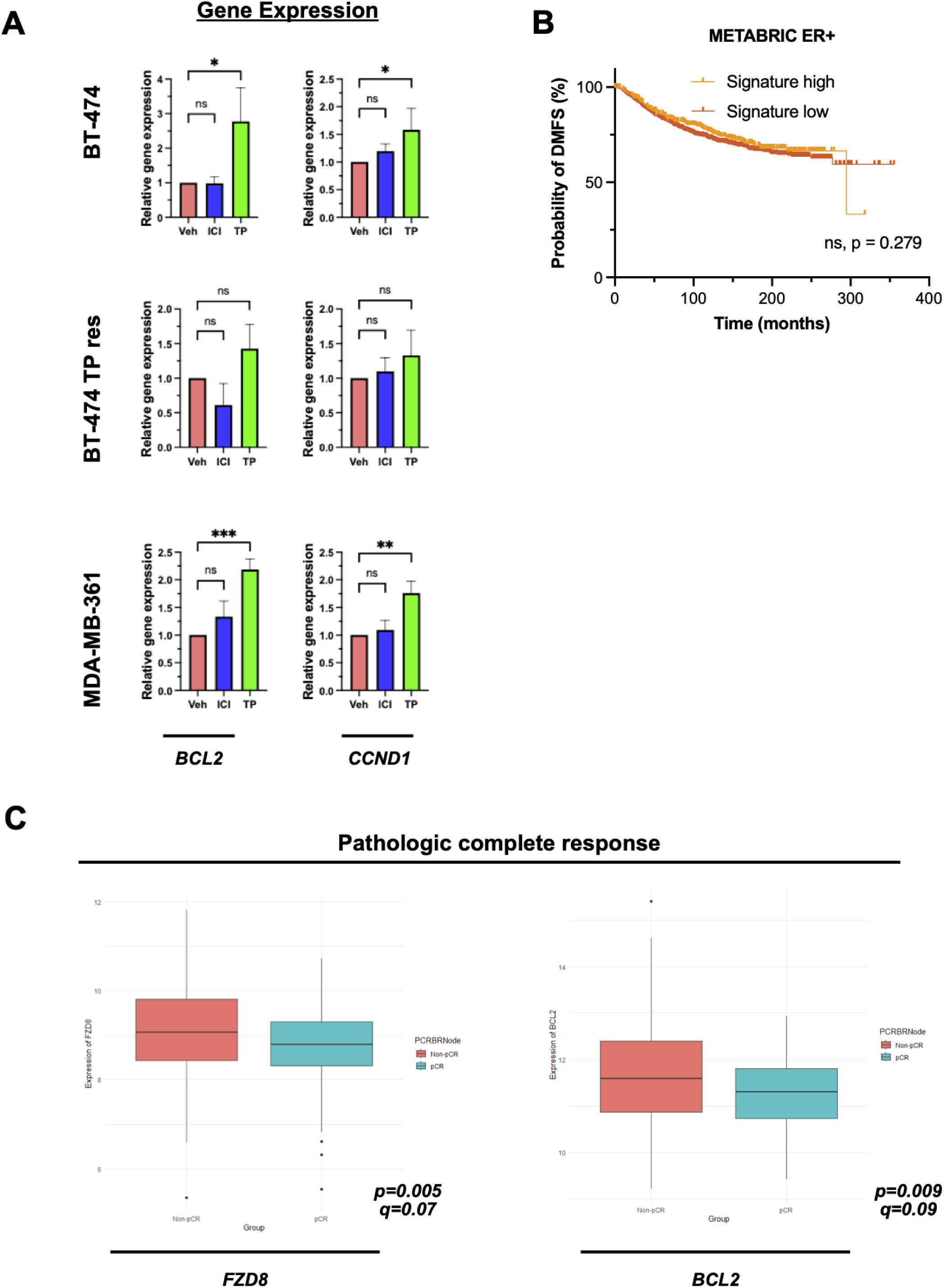
Expression of genes identified by ER CUT&RUN in ER+/HER2+ cells. (A) RNA expression of *BCL2* and *CCND1* in BT-474 parental, BT-474 TP-resistant, and MDA-MB-361 cells treated with ICI and TP for 24 hours. Data from 3 independent biological replicates is presented as the mean ± SD and analyzed by one-way ANOVA with Dunnett’s multiple comparisons test. (B) Distant metastasis-free survival (DMFS) of ER+ breast cancer patients in the METABRIC cohort (n=1363), separated by high (orange, n=379) or low (red, n=984) median expression of the same ER- bound 39-gene signature derived from the top 5 enriched pathways in Fig. 5C and Table S4. A log-rank Mantel-Cox test was used. (C) Expression of *FZD8* or *BCL2* in breast tumors of patients with HER2+ breast cancer from the overall cohort of the NSABP B-41 clinical trial (n=245; ER+ = 141, ER- = 104), stratified by pCR status. \**p* < 0.05, \*\**p* < 0.01, and \*\*\**p* < 0.001; ns, not significant.

**Table S4.** Predicted ER target genes in TP-treated BT-474 cells.

| Wnt signaling pathway | Melanogenesis | NOD-like receptor signaling pathway | Mitophagy - animal | AGE-RAGE signaling pathway in diabetic complications | All - Duplicates Removed |
| --- | --- | --- | --- | --- | --- |
| TBL1X | FZD9 | NLRP12 | CCZ1B | COL4A3 | TBL1X |
| FZD9 | FZD8 | RELA | RELA | RELA | FZD9 |
| FZD8 | WNT11 | NEK7 | TOMM20 | PIM1 | FZD8 |
| SIRT1 | PRKCB | IKBKE | MAP1LC3C | CCND1 | SIRT1 |
| CSNK2A3 | FZD7 | MAP1LC3C | CSNK2A3 | COL4A2 | CSNK2A3 |
| CCND1 | RAF1 | JAK1 | RRAS2 | PRKCB | CCND1 |
| WNT11 | DVL3 | GBP3 |  | SMAD4 | WNT11 |
| WIF1 | MITF | DNM1L |  | BCL2 | WIF1 |
| PRKCB | LEF1 | PYDC1 |  |  | PRKCB |
| SMAD4 | CAMK2A | BCL2 |  |  | SMAD4 |
| FZD7 | ADCY2 | TANK |  |  | FZD7 |
| CSNK1E | EDN1 |  |  |  | CSNK1E |
| RUVBL1 |  |  |  |  | RUVBL1 |
| DVL3 |  |  |  |  | DVL3 |
| DKK2 |  |  |  |  | DKK2 |
| LEF1 |  |  |  |  | LEF1 |
| CTNND2 |  |  |  |  | CTNND2 |
| CAMK2A |  |  |  |  | CAMK2A |
|  |  |  |  |  | RAF1 |
|  |  |  |  |  | MITF |
|  |  |  |  |  | ADCY2 |
|  |  |  |  |  | EDN1 |
|  |  |  |  |  | NLRP12 |
|  |  |  |  |  | RELA |
|  |  |  |  |  | NEK7 |
|  |  |  |  |  | IKBKE |
|  |  |  |  |  | MAP1LC3C |
|  |  |  |  |  | JAK1 |
|  |  |  |  |  | GBP3 |
|  |  |  |  |  | DNM1L |
|  |  |  |  |  | PYDC1 |
|  |  |  |  |  | BCL2 |
|  |  |  |  |  | TANK |
|  |  |  |  |  | CCZ1B |
|  |  |  |  |  | TOMM20 |
|  |  |  |  |  | RRAS2 |
|  |  |  |  |  | COL4A3 |
|  |  |  |  |  | PIM1 |
|  |  |  |  |  | COL4A2 |

## Acknowledgments

The authors would like to thank the patients that donated samples for the construction of the breast tumor tissue microarray, the members of the Riggins laboratory for their support, the Sreejith Nair laboratory (Georgetown University) for their support with ChIP experiments, and the Matthew Sikora laboratory (University of Colorado Anschutz) for their insights.

## Author Contributions

S.T., S.B., S.R., S.M.S., and R.B.R. designed the experiments. S.T. performed most of the experiments, analyzed data from most of the experiments, prepared figures and wrote the manuscript. S.B. contributed to the conceptualization of the presented work, designed, performed and analyzed the immunofluorescence experiments, and edited the manuscript and prepared figures. L.J. analyzed and created figures for the CUT&RUN experiment. M.D.M. and A.T.M. analyzed and created figures for the ChIP-seq experiment. S.R. designed, wrote code for, and performed the analysis of the HER2 TMA data. G.T. performed analysis of NSABP B-41 RNA sequencing data. R.R. and D.K.K. acquired the images for the immunofluorescence experiments. R.B.R. oversaw the study, wrote, prepared figures, and reviewed and edited the manuscript. All authors read, revised, and approved the manuscript.

## Abbreviations

ADCC: Antibody-dependent cellular cytotoxicity
BETA: Binding and expression target analysis
ChIP-qPCR: Chromatin immunoprecipitation and quantitative polymerase chain reaction
ChIP-seq: Chromatin immunoprecipitation sequencing
CUT&RUN: Cleavage under targets and release using nuclease
DMFS: Distant metastasis-free survival
E2 17β: estradiol
EGF: Epidermal growth factor
EGFR: Epidermal growth factor receptor
ER: Estrogen receptor alpha
ERE: Estrogen response element
GO: Gene ontology
GSEA: Gene set enrichment analysis
HER2: Human epidermal growth factor receptor 2
HR: Hormone receptor
IHC: Immunohistochemistry
ICI: Fulvestrant
pCR: Pathologic complete response
PFS: Progression-free survival
pHER2: Phospho-HER2
PTM: Post-translational modification
qPCR: Quantitative polymerase chain reaction
RAGE: Receptor for advanced glycation endproducts
TKI: Tyrosine kinase inhibitor
TMA: Tissue microarray
TME: Tumor microenvironment
TP: Trastuzumab and pertuzumab
Tuc: Tucatinib

## References

1. Siegel RL, Kratzer TB, Wagle NS, Sung H, Jemal A. Cancer statistics, 2026. CA Cancer J Clin. 2026 Jan 1;76(1):e70043. doi:10.3322/caac.70043

2. Allison KH, Hammond MEH, Dowsett M, McKernin SE, Carey LA, Fitzgibbons PL, et al. Estrogen and Progesterone Receptor Testing in Breast Cancer: ASCO/CAP Guideline Update. J Clin Oncol. 2020 Apr 20;38(12):1346–66. doi:10.1200/JCO.19.02309

3. Wolff AC, Somerfield MR, Dowsett M, Hammond MEH, Hayes DF, McShane LM, et al. Human Epidermal Growth Factor Receptor 2 Testing in Breast Cancer: ASCO–College of American Pathologists Guideline Update. J Clin Oncol. 2023 Aug;41(22):3867–72. doi:10.1200/JCO.22.02864

4. Corti C, Batra-Sharma H, Kelsten M, Shatsky RA, Garrido-Castro AC, Gradishar WJ. Systemic Therapy in Breast Cancer. Am Soc Clin Oncol Educ Book. 2024 Jan 7;44(3):e432442. doi:10.1200/EDBK_432442

5. Korde LA, Somerfield MR, Carey LA, Crews JR, Denduluri N, Hwang ES, et al. Neoadjuvant Chemotherapy, Endocrine Therapy, and Targeted Therapy for Breast Cancer: ASCO Guideline. J Clin Oncol. 2021 May;39(13):1485–505. doi:10.1200/JCO.20.03399

6. Bar M, Widłak W, Vydra N. Targeting ERα Coregulator Networks to Overcome Endocrine Resistance in ER+ Breast Cancer. Int J Cancer. 2026 May 4. doi:10.1002/ijc.70522 PubMed PMID: 42083279.

7. Swain SM, Shastry M, Hamilton E. Targeting HER2-positive breast cancer: advances and future directions. Nat Rev Drug Discov. 2023 Feb;22(2):101–26. doi:10.1038/s41573-022-00579-0

8. Pegram M, Jackisch C, Johnston SRD. Estrogen/HER2 receptor crosstalk in breast cancer: combination therapies to improve outcomes for patients with hormone receptor-positive/HER2- positive breast cancer. Npj Breast Cancer. 2023 May 31;9(1):45. doi:10.1038/s41523-023-00533-2

9. Tommasi C, Airò G, Pratticò F, Testi I, Corianò M, Pellegrino B, et al. Hormone Receptor-Positive/HER2-Positive Breast Cancer: Hormone Therapy and Anti-HER2 Treatment: An Update on Treatment Strategies. J Clin Med. 2024 Mar 24;13(7):1873. doi:10.3390/jcm13071873 PubMed PMID: 38610638; PubMed Central PMCID: PMC11012464.

10. Giordano SH, Franzoi MAB, Temin S, Anders CK, Chandarlapaty S, Crews JR, et al. Systemic Therapy for Advanced Human Epidermal Growth Factor Receptor 2–Positive Breast Cancer: ASCO Guideline Update. J Clin Oncol. 2022 Aug 10;40(23):2612–35. doi:10.1200/JCO.22.00519

11. Bahnassy S, Stires H, Jin L, Tam S, Mobin D, Balachandran M, et al. Unraveling Vulnerabilities in Endocrine Therapy-Resistant HER2+/ER+ Breast Cancer. Endocrinology. 2023 Dec 1;164(12):bqad159. doi:10.1210/endocr/bqad159

12. Liu S, Yu M, Mou E, Wang M, Liu S, Xia L, et al. The optimal neoadjuvant treatment strategy for HR+/HER2 + breast cancer: a network meta-analysis. Sci Rep. 2025 Jan 3;15(1):713. doi:10.1038/s41598-024-84039-2 PubMed PMID: 39753653; PubMed Central PMCID: PMC11699132.

13. Chen H, Gui X, Zhou Z, Su F, Gong C, Li S, et al. Distinct ER and PR expression patterns significantly affect the clinical outcomes of early HER2-positive breast cancer: A real-world analysis of 871 patients treated with neoadjuvant therapy. The Breast. 2024 Jun 1;75. doi:10.1016/j.breast.2024.103733 PubMed PMID: 38615482.

14. Robidoux A, Tang G, Rastogi P, Geyer CE, Azar CA, Atkins JN, et al. Lapatinib as a component of neoadjuvant therapy for HER2-positive operable breast cancer (NSABP protocol B-41): an open-label, randomised phase 3 trial. Lancet Oncol. 2013 Nov 1;14(12):1183–92. doi:10.1016/S1470-2045(13)70411-X

15. Baselga J, Bradbury I, Eidtmann H, Di Cosimo S, de Azambuja E, Aura C, et al. Lapatinib with trastuzumab for HER2-positive early breast cancer (NeoALTTO): a randomised, open-label, multicentre, phase 3 trial. The Lancet. 2012 Feb 18;379(9816):633–40. doi:10.1016/S0140-6736(11)61847-3

16. Gianni L, Pienkowski T, Im YH, Roman L, Tseng LM, Liu MC, et al. Efficacy and safety of neoadjuvant pertuzumab and trastuzumab in women with locally advanced, inflammatory, or early HER2-positive breast cancer (NeoSphere): a randomised multicentre, open-label, phase 2 trial. Lancet Oncol. 2012 Jan 1;13(1):25–32. doi:10.1016/S1470-2045(11)70336-9 PubMed PMID: 22153890.

17. Schneeweiss A, Chia S, Hickish T, Harvey V, Eniu A, Hegg R, et al. Pertuzumab plus trastuzumab in combination with standard neoadjuvant anthracycline-containing and anthracycline-free chemotherapy regimens in patients with HER2-positive early breast cancer: a randomized phase II cardiac safety study (TRYPHAENA). Ann Oncol Off J Eur Soc Med Oncol. 2013 Sep;24(9):2278–84. doi:10.1093/annonc/mdt182 PubMed PMID: 23704196.

18. Carey LA, Berry DA, Cirrincione CT, Barry WT, Pitcher BN, Harris LN, et al. Molecular Heterogeneity and Response to Neoadjuvant Human Epidermal Growth Factor Receptor 2 Targeting in CALGB 40601, a Randomized Phase III Trial of Paclitaxel Plus Trastuzumab With or Without Lapatinib. J Clin Oncol. 2016 Feb 20;34(6):542–9. doi:10.1200/JCO.2015.62.1268 PubMed PMID: 26527775; PubMed Central PMCID: PMC4980567.

19. Martínez-Sáez O, Prat A. Current and Future Management of HER2-Positive Metastatic Breast Cancer. JCO Oncol Pract. 2021 Oct;17(10):594–604. doi:10.1200/OP.21.00172

20. Callari M, Dugo M, Barreca M, Győrffy B, Galbardi B, Vigano L, et al. Determinants of response and molecular dynamics in HER2+ER+ breast cancers from the NA-PHER2 trial receiving HER2- targeted and endocrine therapies. Nat Commun. 2025 Mar 4;16:2195. doi:10.1038/s41467-025-57293-9 PubMed PMID: 40038334; PubMed Central PMCID: PMC11880565.

21. Li KC, Girardi E, Kartnig F, Grosche S, Pemovska T, Bigenzahn JW, et al. Cell-surface SLC nucleoside transporters and purine levels modulate BRD4-dependent chromatin states. Nat Metab. 2021 May;3(5):651–64. doi:10.1038/s42255-021-00386-8

22. Schindelin J, Arganda-Carreras I, Frise E, Kaynig V, Longair M, Pietzsch T, et al. Fiji: an open-source platform for biological-image analysis. Nat Methods. 2012 Jul;9(7):676–82. doi:10.1038/nmeth.2019

23. Bentley DR, Balasubramanian S, Swerdlow HP, Smith GP, Milton J, Brown CG, et al. Accurate Whole Human Genome Sequencing using Reversible Terminator Chemistry. Nature. 2008 Nov 6;456(7218):53–9. doi:10.1038/nature07517 PubMed PMID: 18987734; PubMed Central PMCID: PMC2581791.

24. Li H, Durbin R. Fast and accurate short read alignment with Burrows–Wheeler transform. Bioinformatics. 2009 Jul 15;25(14):1754–60. doi:10.1093/bioinformatics/btp324 PubMed PMID: 19451168; PubMed Central PMCID: PMC2705234.

25. Gaspar JM. Improved peak-calling with MACS2 [Internet]. bioRxiv; 2018 [cited 2026 May 17]. p. 496521. Available from: https://www.biorxiv.org/content/10.1101/496521v1 doi:10.1101/496521

26. Quinlan AR, Hall IM. BEDTools: a flexible suite of utilities for comparing genomic features. Bioinformatics. 2010 Mar 15;26(6):841–2. doi:10.1093/bioinformatics/btq033 PubMed PMID: 20110278; PubMed Central PMCID: PMC2832824.

27. Ramírez F, Ryan DP, Grüning B, Bhardwaj V, Kilpert F, Richter AS, et al. deepTools2: a next generation web server for deep-sequencing data analysis. Nucleic Acids Res. 2016 Jul 8;44(W1):W160–5. doi:10.1093/nar/gkw257

28. Wang S, Sun H, Ma J, Zang C, Wang C, Wang J, et al. Target analysis by integration of transcriptome and ChIP-seq data with BETA. Nat Protoc. 2013 Dec;8(12):2502–15. doi:10.1038/nprot.2013.150 PubMed PMID: 24263090; PubMed Central PMCID: PMC4135175.

29. Guan J, Zhou W, Hafner M, Blake RA, Chalouni C, Chen IP, et al. Therapeutic Ligands Antagonize Estrogen Receptor Function by Impairing Its Mobility. Cell. 2019 Aug 8;178(4):949–963.e18. doi:10.1016/j.cell.2019.06.026 PubMed PMID: 31353221.

30. Subramanian A, Tamayo P, Mootha VK, Mukherjee S, Ebert BL, Gillette MA, et al. Gene set enrichment analysis: A knowledge-based approach for interpreting genome-wide expression profiles. Proc Natl Acad Sci. 2005 Oct 25;102(43):15545–50. doi:10.1073/pnas.0506580102

31. Mootha VK, Lindgren CM, Eriksson KF, Subramanian A, Sihag S, Lehar J, et al. PGC-1α- responsive genes involved in oxidative phosphorylation are coordinately downregulated in human diabetes. Nat Genet. 2003 Jul;34(3):267–73. doi:10.1038/ng1180

32. Marunde MR, Fuchs HA, Burg JM, Popova IK, Vaidya A, Hall NW, et al. Nucleosome conformation dictates the histone code. Narlikar GJ, Struhl K, editors. eLife. 2024 Feb 6;13:e78866. doi:10.7554/eLife.78866

33. Yusufova N, Kloetgen A, Teater M, Osunsade A, Camarillo JM, Chin CR, et al. Histone H1 loss drives lymphoma by disrupting 3D chromatin architecture. Nature. 2021 Jan;589(7841):299–305. doi:10.1038/s41586-020-3017-y PubMed PMID: 33299181; PubMed Central PMCID: PMC7855728.

34. Skene PJ, Henikoff JG, Henikoff S. Targeted in situ genome-wide profiling with high efficiency for low cell numbers. Nat Protoc. 2018 May;13(5):1006–19. doi:10.1038/nprot.2018.015

35. Robinson JT, Thorvaldsdóttir H, Winckler W, Guttman M, Lander ES, Getz G, et al. Integrative Genomics Viewer. Nat Biotechnol. 2011 Jan;29(1):24–6. doi:10.1038/nbt.1754 PubMed PMID: 21221095; PubMed Central PMCID: PMC3346182.

36. Fernandez AI, Geng X, Chaldekas K, Harris B, Duttargi A, Berry VL, et al. The orphan nuclear receptor estrogen-related receptor beta (ERRβ) in triple-negative breast cancer. Breast Cancer Res Treat. 2020 Feb;179(3):585–604. doi:10.1007/s10549-019-05485-5 PubMed PMID: 31741180; PubMed Central PMCID: PMC7153462.

37. Olukoya AO, Stires H, Bahnassy S, Persaud S, Guerra Y, Ranjit S, et al. Riluzole Suppresses Growth and Enhances Response to Endocrine Therapy in ER+ Breast Cancer. J Endocr Soc. 2023 Sep 15;7(10):bvad117. doi:10.1210/jendso/bvad117 PubMed PMID: 37766843; PubMed Central PMCID: PMC10521904.

38. Bahnassy S, Young TA, Abalum TC, Pope EA, Rivera AT, Fernandez AI, et al. Glutamate-Handling Proteins Associate with Adverse Clinicopathologic Features and Comorbidities in Invasive Lobular Carcinoma [Internet]. bioRxiv; 2026 [cited 2026 May 17]. p. 2024.09.29.615681. Available from: https://www.biorxiv.org/content/10.1101/2024.09.29.615681v2doi:10.1101/2024.09.29.615681

39. Curtis C, Shah SP, Chin SF, Turashvili G, Rueda OM, Dunning MJ, et al. The genomic and transcriptomic architecture of 2,000 breast tumours reveals novel subgroups. Nature. 2012 Apr 18;486(7403):346–52. doi:10.1038/nature10983 PubMed PMID: 22522925; PubMed Central PMCID: PMC3440846.

40. Pereira B, Chin SF, Rueda OM, Vollan HKM, Provenzano E, Bardwell HA, et al. The somatic mutation profiles of 2,433 breast cancers refines their genomic and transcriptomic landscapes. Nat Commun. 2016 May 10;7:11479. doi:10.1038/ncomms11479 PubMed PMID: 27161491; PubMed Central PMCID: PMC4866047.

41. Rueda OM, Sammut SJ, Seoane JA, Chin SF, Caswell-Jin JL, Callari M, et al. Dynamics of breast-cancer relapse reveal late-recurring ER-positive genomic subgroups. Nature. 2019 Mar;567(7748):399–404. doi:10.1038/s41586-019-1007-8 PubMed PMID: 30867590; PubMed Central PMCID: PMC6647838.

42. Győrffy B. Pharmacologically actionable transcriptomic signatures across immune infiltration subtypes in the KM-plotter breast cancer cohort. Adv Transl Res. 2026 Jun 26;1(aop). doi:10.1556/1661.2026.00008

43. Cerami E, Gao J, Dogrusoz U, Gross BE, Sumer SO, Aksoy BA, et al. The cBio cancer genomics portal: an open platform for exploring multidimensional cancer genomics data. Cancer Discov. 2012 May;2(5):401–4. doi:10.1158/2159-8290.CD-12-0095 PubMed PMID: 22588877; PubMed Central PMCID: PMC3956037.

44. Gao J, Aksoy BA, Dogrusoz U, Dresdner G, Gross B, Sumer SO, et al. Integrative analysis of complex cancer genomics and clinical profiles using the cBioPortal. Sci Signal. 2013 Apr 2;6(269):pl1. doi:10.1126/scisignal.2004088 PubMed PMID: 23550210; PubMed Central PMCID: PMC4160307.

45. de Bruijn I, Kundra R, Mastrogiacomo B, Tran TN, Sikina L, Mazor T, et al. Analysis and Visualization of Longitudinal Genomic and Clinical Data from the AACR Project GENIE Biopharma Collaborative in cBioPortal. Cancer Res. 2023 Dec 1;83(23):3861–7. doi:10.1158/0008-5472.CAN-23-0816 PubMed PMID: 37668528; PubMed Central PMCID: PMC10690089.

46. Fekete JT, Győrffy B. ROCplot.org: Validating predictive biomarkers of chemotherapy/hormonal therapy/anti-HER2 therapy using transcriptomic data of 3,104 breast cancer patients. Int J Cancer. 2019;145(11):3140–51. doi:10.1002/ijc.32369

47. Fernandez-Martinez A, Rediti M, Tang G, Pascual T, Hoadley KA, Venet D, et al. Tumor Intrinsic Subtypes and Gene Expression Signatures in Early-Stage ERBB2/HER2-Positive Breast Cancer: A Pooled Analysis of CALGB 40601, NeoALTTO, and NSABP B-41 Trials. JAMA Oncol. 2024 May 1;10(5):603–11. doi:10.1001/jamaoncol.2023.7304 PubMed PMID: 38546612; PubMed Central PMCID: PMC10979363.

48. Ju J, Du F, Gao SL, Si YR, Hu NL, Liu DX, et al. Combined analysis of receptor expression reflects inter-and intra-tumor heterogeneity in HR+/HER2+ breast cancer. Breast Cancer Res Treat. 2022 Jul 1;194(2):221–30. doi:10.1007/s10549-022-06629-w

49. Masuda S, Nitta H, Kelly BD, Zhang W, Farrell M, Dennis E. Intratumoral Estrogen Receptor Heterogeneity of Expression in Human Epidermal Growth Factor Receptor 2–Positive Breast Cancer as Evaluated by a Brightfield Multiplex Assay. J Histochem Cytochem. 2019 Aug;67(8):563–74. doi:10.1369/0022155419856862 PubMed PMID: 31184528; PubMed Central PMCID: PMC6669858.

50. Diermeier S, Horváth G, Knuechel-Clarke R, Hofstaedter F, Szöllősi J, Brockhoff G. Epidermal growth factor receptor coexpression modulates susceptibility to Herceptin in HER2/neu overexpressing breast cancer cells via specific erbB-receptor interaction and activation. Exp Cell Res. 2005 Apr 1;304(2):604–19. doi:10.1016/j.yexcr.2004.12.008

51. Bon G, Pizzuti L, Laquintana V, Loria R, Porru M, Marchiò C, et al. Loss of HER2 and decreased T-DM1 efficacy in HER2 positive advanced breast cancer treated with dual HER2 blockade: the SePHER Study. J Exp Clin Cancer Res. 2020 Dec 10;39(1):279. doi:10.1186/s13046-020-01797-3

52. Wang L, Paudel BB, McKnight RA, Janes KA. Nucleocytoplasmic transport of active HER2 causes fractional escape from the DCIS-like state. Nat Commun. 2023 Apr 13;14(1):2110. doi:10.1038/s41467-023-37914-x

53. Austin CD, De MAM, Pisacane PI, van DSM, Eigenbrot C, Sliwkowski MX, et al. Endocytosis and Sorting of ErbB2 and the Site of Action of Cancer Therapeutics Trastuzumab and Geldanamycin. Mol Biol Cell. 2004 Dec;15(12):5268–82. doi:10.1091/mbc.e04-07-0591

54. Pedersen MW, Jacobsen HJ, Koefoed K, Dahlman A, Kjær I, Poulsen TT, et al. Targeting Three Distinct HER2 Domains with a Recombinant Antibody Mixture Overcomes Trastuzumab Resistance. Mol Cancer Ther. 2015 Mar 9;14(3):669–80. doi:10.1158/1535-7163.MCT-14-0697

55. Redmond AM, Omarjee S, Chernukhin I, Le Romancer M, Carroll JS. Analysis of HER2 genomic binding in breast cancer cells identifies a global role in direct gene regulation. PLoS ONE. 2019 Nov 20;14(11):e0225180. doi:10.1371/journal.pone.0225180 PubMed PMID:1747426; PubMed Central PMCID: PMC6867699.

56. Duan N, Hua Y, Yan X, He Y, Zeng T, Gong J, et al. Unveiling Alterations of Epigenetic Modifications and Chromatin Architecture Leading to Lipid Metabolic Reprogramming during the Evolutionary Trastuzumab Adaptation of HER2-Positive Breast Cancer. Adv Sci. 2024 Mar 9;11(18):2309424. doi:10.1002/advs.202309424 PubMed PMID: 38460162; PubMed Central PMCID: PMC11095153.

57. Bao Y, Oguz G, Lee WC, Lee PL, Ghosh K, Li J, et al. EZH2-mediated PP2A inactivation confers resistance to HER2-targeted breast cancer therapy. Nat Commun. 2020 Nov 18;11:5878. doi:10.1038/s41467-020-19704-x PubMed PMID: 33208750; PubMed Central PMCID: PMC7674491.

58. Wang YC, Morrison G, Gillihan R, Guo J, Ward RM, Fu X, et al. Different mechanisms for resistance to trastuzumab versus lapatinib in HER2- positive breast cancers -- role of estrogen receptor and HER2 reactivation. Breast Cancer Res BCR. 2011;13(6):R121. doi:10.1186/bcr3067 PubMed PMID: 22123186; PubMed Central PMCID: PMC3326563.

59. Parida PK, Marquez-Palencia M, Nair V, Kaushik AK, Kim K, Sudderth J, et al. Metabolic diversity within breast cancer brain-tropic cells determines metastatic fitness. Cell Metab. 2022 Jan 4;34(1):90–105.e7. doi:10.1016/j.cmet.2021.12.001 PubMed PMID: 34986341; PubMed Central PMCID: PMC9307073.

60. Rojas A, Lindner C, Schneider I, Gonzalez I, Uribarri J. The RAGE Axis: A Relevant Inflammatory Hub in Human Diseases. Biomolecules. 2024 Apr;14(4):412. doi:10.3390/biom14040412

61. Dai X, Hou Y, Deng T, Lin G, Cao Y, Yu G, et al. A specific RAGE-binding peptide inhibits triple negative breast cancer growth through blocking of Erk1/2/NF-κB pathway. Eur J Pharmacol. 2023 Sep 5;954:175861. doi:10.1016/j.ejphar.2023.175861

62. Eeckhoute J, Carroll JS, Geistlinger TR, Torres-Arzayus MI, Brown M. A cell-type-specific transcriptional network required for estrogen regulation of cyclin D1 and cell cycle progression in breast cancer. Genes Dev. 2006 Sep 15;20(18):2513–26. doi:10.1101/gad.1446006 PubMed PMID: 16980581; PubMed Central PMCID: PMC1578675.

63. Perillo B, Sasso A, Abbondanza C, Palumbo G. 17beta-estradiol inhibits apoptosis in MCF-7 cells, inducing bcl-2 expression via two estrogen-responsive elements present in the coding sequence. Mol Cell Biol. 2000 Apr;20(8):2890–901. doi:10.1128/MCB.20.8.2890-2901.2000 PubMed PMID: 10733592; PubMed Central PMCID: PMC85519.

64. Swain SM, Tang G, Brauer HA, Goerlitz DS, Lucas PC, Robidoux A, et al. NSABP B-41, a Randomized Neoadjuvant Trial: Genes and Signatures Associated with Pathologic Complete Response. Clin Cancer Res Off J Am Assoc Cancer Res. 2020 Aug 15;26(16):4233–41. doi:10.1158/1078-0432.CCR-20-0152 PubMed PMID: 32371537; PubMed Central PMCID: PMC7724952.

65. Metzger O, Mandrekar S, Goel S, Gligorov J, Lim E, Ciruelos E, et al. Palbociclib for Hormone-Receptor–Positive, HER2-Positive Advanced Breast Cancer. N Engl J Med. 2026 Jan 28;394(5):451–62. doi:10.1056/NEJMoa2511218

66. Liu C, Sun L, Niu N, Hou P, Chen G, Wang H, et al. Molecular classification of hormone receptor-positive /HER2-positive breast cancer reveals potential neoadjuvant therapeutic strategies. Signal Transduct Target Ther. 2025 Mar 26;10(1):97. doi:10.1038/s41392-025-02181-3

67. Dillon MF, Stafford AT, Kelly G, Redmond AM, McIlroy M, Crotty TB, et al. Cyclooxygenase-2 predicts adverse effects of tamoxifen: a possible mechanism of role for nuclear HER2 in breast cancer patients. Endocr Relat Cancer. 2008 Sep;15(3):745–53. doi:10.1677/ERC-08-0009 PubMed PMID: 18469157.

68. Wang SC, Lien HC, Xia W, Chen IF, Lo HW, Wang Z, et al. Binding at and transactivation of the COX-2 promoter by nuclear tyrosine kinase receptor ErbB-2. Cancer Cell. 2004 Sep 1;6(3):251– 61. doi:10.1016/j.ccr.2004.07.012 PubMed PMID: 15380516.

69. Li LY, Chen H, Hsieh YH, Wang YN, Chu HJ, Chen YH, et al. Nuclear ErbB-2 Enhances Translation and Cell Growth by activating transcription of rRNA genes. Cancer Res. 2011 Jun 15;71(12):4269–79. doi:10.1158/0008-5472.CAN-10-3504 PubMed PMID: 21555369; PubMed Central PMCID: PMC3117049.

70. Pereira PMR, Sharma SK, Carter LM, Edwards KJ, Pourat J, Ragupathi A, et al. Caveolin-1 mediates cellular distribution of HER2 and affects trastuzumab binding and therapeutic efficacy. Nat Commun. 2018 Dec 3;9(1):5137. doi:10.1038/s41467-018-07608-w

71. Mishra A, Hourigan D, Lindsay AJ. Inhibition of the endosomal recycling pathway downregulates HER2 activation and overcomes resistance to tyrosine kinase inhibitors in HER2-positive breast cancer. Cancer Lett. 2022 Mar 31;529:153–67. doi:10.1016/j.canlet.2022.01.003

72. Murad R, Avanes A, Ma X, Geng S, Mortazavi A, Momand J. Transcriptome and chromatin landscape changes associated with trastuzumab resistance in HER2+ breast cancer cells. Gene. 2021 Oct 5;799:145808. doi:10.1016/j.gene.2021.145808

73. Kokot A, Gadakh S, Saha I, Gajda E, Łaźniewski M, Rakshit S, et al. Unveiling the Molecular Mechanism of Trastuzumab Resistance in SKBR3 and BT474 Cell Lines for HER2 Positive Breast Cancer. Curr Issues Mol Biol. 2024 Mar 21;46(3):2713–40. doi:10.3390/cimb46030171 PubMed PMID: 38534787; PubMed Central PMCID: PMC10969458.

74. Mandó P, Rivero SG, Rizzo MM, Pinkasz M, Levy EM. Targeting ADCC: A different approach to HER2 breast cancer in the immunotherapy era. Breast Off J Eur Soc Mastology. 2021 Aug 19;60:15–25. doi:10.1016/j.breast.2021.08.007 PubMed PMID: 34454323; PubMed Central PMCID: PMC8399304.

75. Murthy RK, Loi S, Okines A, Paplomata E, Hamilton E, Hurvitz SA, et al. Tucatinib, Trastuzumab, and Capecitabine for HER2-Positive Metastatic Breast Cancer. N Engl J Med. 2020 Feb 13;382(7):597–609. doi:10.1056/NEJMoa1914609

76. Lupien M, Meyer CA, Bailey ST, Eeckhoute J, Cook J, Westerling T, et al. Growth factor stimulation induces a distinct ERα cistrome underlying breast cancer endocrine resistance. Genes Dev. 2010 Oct 1;24(19):2219–27. doi:10.1101/gad.1944810 PubMed PMID: 20889718.

77. Luque M, Morales-Gallego M, Sanz-Álvarez M, Arguiñano C, Rangel Y, Ramírez-Merino N, et al. Cancer-associated fibroblast-derived protein S100-A11 influences the response to anti-HER2 therapies in HER2-positive breast cancer. Neoplasia N Y N. 2026 May 19;78:101318. doi:10.1016/j.neo.2026.101318 PubMed PMID: 42155177; PubMed Central PMCID: PMC13213234.

78. Triulzi T, Forte L, Regondi V, Di Modica M, Ghirelli C, Carcangiu ML, et al. HER2 signaling regulates the tumor immune microenvironment and trastuzumab efficacy. Oncoimmunology. 2018 Sep 21;8(1):e1512942. doi:10.1080/2162402X.2018.1512942 PubMed PMID: 30546951; PubMed Central PMCID: PMC6287794.

79. Kamashev D, Shaban N, Suntsova M, Raevskiy M, Efimov V, Moisseev A, et al. Human Blood Serum Inhibits Ductal Carcinoma Cells BT474 Growth and Modulates Effect of HER2 Inhibition. Biomedicines. 2022 Aug 8;10(8):1914. doi:10.3390/biomedicines10081914 PubMed PMID: 36009461; PubMed Central PMCID: PMC9405390.

80. Pietilä M, Sahgal P, Peuhu E, Jäntti NZ, Paatero I, Närvä E, et al. SORLA regulates endosomal trafficking and oncogenic fitness of HER2. Nat Commun. 2019 May 28;10(1):2340. doi:10.1038/s41467-019-10275-0

81. Hendriks BS, Wiley HS, Lauffenburger D. HER2-Mediated Effects on EGFR Endosomal Sorting: Analysis of Biophysical Mechanisms. Biophys J. 2003 Oct;85(4):2732–45. doi:10.1016/s0006-3495(03)74696-7 PubMed PMID: 14507736; PubMed Central PMCID: PMC1303497.

82. Bai X, Sun P, Wang X, Long C, Liao S, Dang S, et al. Structure and dynamics of the EGFR/HER2 heterodimer. Cell Discov. 2023 Feb 13;9(1):18. doi:10.1038/s41421-023-00523-5

83. Akar U, Chaves-Reyez A, Barria M, Tari A, Sanguino A, Kondo Y, et al. Silencing of Bcl-2 expression by small interfering RNA induces autophagic cell death in MCF-7 breast cancer cells. Autophagy. 2008 Jul 1;4(5):669–79. doi:10.4161/auto.6083 PubMed PMID: 18424910.

84. Santolla MF, Talia M, Maggiolini M. S100A4 Is Involved in Stimulatory Effects Elicited by the FGF2/FGFR1 Signaling Pathway in Triple-Negative Breast Cancer (TNBC) Cells. Int J Mol Sci. 2021 Apr 29;22(9):4720. doi:10.3390/ijms22094720 PubMed PMID: 33946884; PubMed Central PMCID: PMC8124532.

